# Cooperative Chikungunya virus membrane fusion and its sub-stoichiometric inhibition by CHK-152 antibody

**DOI:** 10.1101/2020.03.18.996512

**Authors:** Jelle S. Blijleven, Ellen M. Bouma, Mareike K.S. van Duijl, Jolanda M. Smit, Antoine M. van Oijen

## Abstract

Chikungunya virus (CHIKV) presents a major burden on healthcare systems worldwide, but specific treatment remains unavailable. Attachment and fusion of CHIKV to the host cell membrane is mediated by the E1/E2 protein spikes. We used an in vitro single-particle fusion assay to study the effect of the potent, neutralizing antibody CHK-152 on CHIKV binding and fusion. We find that CHK-152 shields the virions, inhibiting interaction with the target membrane and inhibiting fusion. Analysis of the ratio of bound antibodies to epitopes implied that CHIKV fusion is a highly cooperative process. Further, dissociation of the antibody at lower pH results in a finely balanced kinetic competition between inhibition and fusion, suggesting a window of opportunity for the spike proteins to act and mediate fusion even in the presence of antibody.

## Introduction

Chikungunya virus (CHIKV; *Alphavirus* genus, *Togaviridae* family) is a human arthropod-borne virus causing chikungunya fever and potentially long-lasting effects such as joint pain. It has recently greatly expanded its geographic range to encompass most tropical-to-temperate regions of the world (Centers for Disease Control and Prevention (CDC),) and is likely to spread further due to geographic expansion of the mosquito vectors that transmit the virus (Bonizzoni et al., 2013, Reiter, Fontenille & Paupy, 2006, Weaver, Forrester, 2015). No preventive medicine or specific antiviral treatment is available to counter CHIKV infection.

Alphaviruses are enveloped viruses in which the lipid bilayer is derived from the host plasma membrane (Jose, Snyder & Kuhn, 2009). The membrane encapsulates the protein capsid in which the viral genome resides. Two viral proteins, E1 and E2, are anchored in the membrane and arranged in trimers of E1/E2 heterodimers called spikes. The spikes cover the surface in an icosahedral lattice with triangulation *T* = 4, giving rise to 80 spikes, or 240 copies of the E1-E2 heterodimers in total (Voss et al., 2010). The E2 protein facilitates alphavirus binding to cellular receptors (Smith et al., 1995, Ashbrook et al., 2014), and both the E1 and E2 proteins play an important role in the process of membrane fusion.

A critical step in the reproduction cycle of enveloped viruses involves the merger of the viral membrane with the host cellular membrane to deliver the genome to the host cell and start a new cycle of viral replication (reviewed by Harrison (Harrison, 2015)). However, membrane fusion does not occur spontaneously on biological timescales due to high kinetic barriers between the intermediates (Chernomordik, Kozlov, 2008). Enveloped viruses therefore have envelope proteins that catalyze membrane fusion (reviewed by Kielian (Kielian, 2014)), to deliver the viral genome at the right time to the right place in the host cell. Upon attachment of CHIKV to the cell, the virion is taken up into an endosomal compartment, mainly by clathrin-mediated endocytosis (Bernard et al., 2010). Membrane fusion is initiated at the mildly acidic pH of the early endosome (Hoornweg et al., 2016, van Duijl-Richter et al., 2015), triggering the E1-E2 heterodimers to dissociate (Voss et al., 2010, Wahlberg, Boere & Garoff, 1989). The E1 proteins subsequently insert into the endosomal membrane and trimerize to form the functional units of fusion (Wahlberg et al., 1992, Cao, Zhang, 2013). Multiple trimers are thought to be necessary to concertedly bring both membranes together (van Duijl-Richter et al., 2015, Zheng et al., 2011, Gibbons et al., 2004), first leading to a hemifused intermediate where the proximal leaflets have merged, and finally opening a pore to deliver the viral genome into the cellular cytosol.

There is currently no vaccine or treatment available against CHIKV, but several promising antibodies have been isolated and were shown to prevent CHIKV infection (Clayton, 2016). A potent antibody is CHK-152, that was found to protect against CHIKV infection in mouse and non-human primate models (Pal et al., 2013, Pal et al., 2014). Mutational and cryo-EM reconstruction studies showed that it binds to the acid-sensitive region of E2. This region becomes disordered at low pH thereby facilitating exposure of the E1 fusion loop (Voss et al., 2010, Li et al., 2010, Sun et al., 2013).

In this study, we found that CHK-152 strongly interferes with CHIKV membrane interactions both at neutral and low pH. Additionally, in a single-particle fluorescence microscopy assay, fusion of particles that were already docked to the membrane was blocked and slowed down. At pH 6.1 and sub-stoichiometric antibody binding, fusion was efficiently inhibited. This effect was diminished at pH 5 and 4.7 as at these pH values CHK-152 was found to dissociate from the virus particles. We explain the results in a model of CHIKV fusion as mediated by multiple E1 trimers formed from CHK-152-free spikes. The stoichiometry of antibody binding implies a cooperative fusion mechanism, where three to five neighboring E1 trimers are required to mediate membrane fusion.

## Results

To delineate the different mechanistic effects of the CHK-152 antibody on the fusion process we set out to separately characterize membrane binding and fusion. By using a combination of binding assays, we studied the effect of CHK-152 on membrane interactions. These experiments were followed by a single-particle assay with pre-docked particles to directly investigate the effect of CHK-152 on membrane fusion, and to determine the stoichiometry of neutralization.

### CHK-152 shields virions thereby preventing neutral-pH membrane interaction

First, we wanted to determine the effect of CHK-152 on nonspecific membrane interaction at neutral pH. A planar, lipid membrane was formed by using a flow cell constructed on top of a hydrophilic microscope coverslip and introducing liposomes (Floyd et al., 2008). The receptor-free bilayer incorporated DOPC, DOPE, sphingomyelin and cholesterol, the latter two lipids being stimulating and required factors for fusion (van Duijl-Richter et al., 2015, Klimjack, Jeffrey & Kielian, 1994, Nieva et al., 1994, Ahn, Gibbons & Kielian, 2002). CHIKV particles were UV inactivated to render them non-infectious and were labeled with the lipophilic dye R18. After labeling, they were incubated with varying concentrations of CHK-152 antibody and flown into the flow cell to dock to the membrane. After rinsing with buffer, the number of particles sticking to the bilayer was quantified by single-particle fluorescence microscopy (more detail below, in Figure 3 and Methods). Particle counts normalized to the same conditions but in the absence of CHK-152 are shown in Figure 1a on double log scale.

**Figure 1.**
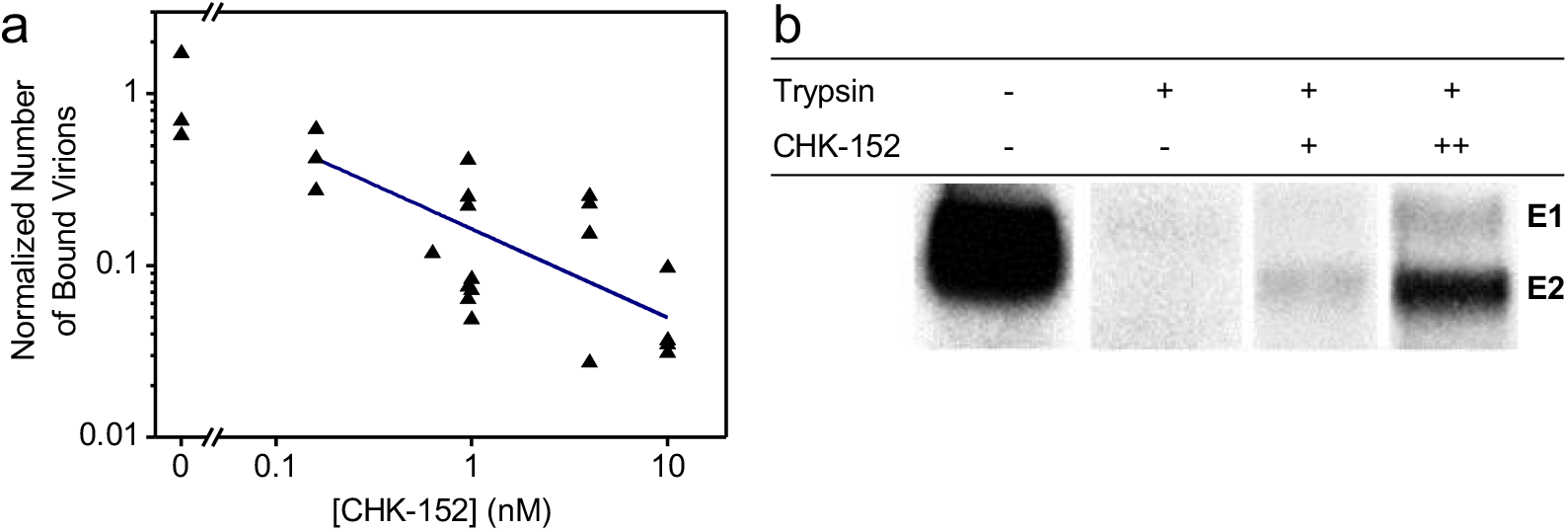
Shielding of virions by CHK-152 at neutral pH. (a) Inhibition of nonspecific binding to a planar membrane. Fluorescently labeled CHIK virions were incubated with CHK-152, flown into a flow cell and docked to a planar membrane (see text). The number of particles binding to the membrane after rinsing the channel was counted and normalized to the mean number of particles in the absence of antibody. Single trials shown on log-log scale (*n* = 25); blue line indicates a power-law fit with power coefficient −0.5±0.2. (b) Shielding of surface proteins from enzymatic cleavage. [35S]-methionine/L-[35S] cysteine labeled CHIKV was incubated with CHK-152 and mixed with liposomes at neutral pH. The mixture was trypsinized for 1 h and subjected to SDS-PAGE analysis. CHK-152 concentration in final volume: +, 0.63 nM CHK-152 in estimated ratio of 13 to virions; ++, 10 nM CHK-152 in ratio of 210 to virions. Representative image out of 3 trials shown.

We found that nonspecific binding reduced with increasing concentration of CHK-152 during the pre-incubation phase, as indicated by the fit of a power function (linear on log-log scale). Interestingly, we also found that CHK-152 shields the E2 surface glycoprotein from enzymatic cleavage by trypsin (Figure 1b). Radiolabeled CHIKV was mixed with liposomes at neutral pH and subjected to trypsin digestion and SDS-page analysis. Trypsin completely digested the E1 and E2 proteins, while pre-incubation with increasing concentrations of CHK-152 protected the E2 protein from trypsin digestion, indicating that E2 proteins were shielded against enzymatic cleavage. Collectively, these results suggest that the CHIKV membrane interaction at neutral pH is reduced due to steric hindrance caused by the CHK-152 antibody.

### CHK-152 blocks interaction with target membranes at low pH

At low pH, the viral fusion proteins undergo conformational changes to support membrane fusion. Antibodies have been described that prevent the conformational changes that are required for membrane fusion or that freeze virus particles in an intermediate stage (Jin et al., 2015, Fox et al., 2015, Kaufmann et al., 2010, Selvarajah et al., 2013, Smith et al., 2015). We described before that CHK-152 abolishes membrane fusion activity at high antibody concentration in a liposomal fusion assay (Pal et al., 2013). There, we investigated the effect of CHK-152 on CHIKV fusion and revealed that both the extent as well as the rate of fusion decreases with increasing antibody concentrations (Pal et al., 2013, van Duijl-Richter, 2016). At 10 nM CHK-152, membrane fusion was almost completely abolished.

To further dissect the role of CHK-152 on membrane fusion, we here determined the low-pH dependent binding properties of the virus to liposomes in the presence or absence of CHK-152, by use of a liposomal co-floatation assay (Figure 2a).

**Figure 2.**
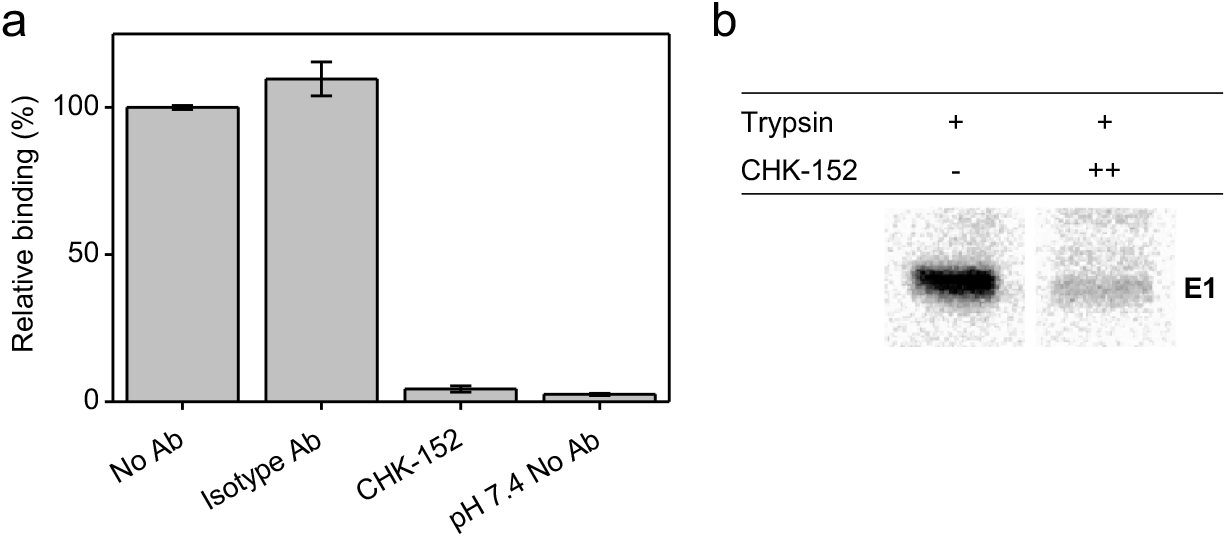
CHK-152 inhibition of target membrane interaction at low pH. (a) Inhibition of E1-liposome interaction at low pH. A fusion experiment was performed, adding radiolabeled CHIKV that was pre-incubated without, or with 10 nM of isotype control or CHK-152 antibodies to liposomes and acidifying the mixture to pH 5.1. After 1 min, the sample was neutralized and added to the bottom of a sucrose gradient and centrifuged. The relative radioactivity in the top fractions, therefore co-floating with the liposomes, was determined in triplicate and is plotted as mean±sem. (b) Inhibition of formation of trypsin-resistant E1 trimer. Radiolabeled CHIKV was incubated with or without CHK-152 for 10 min at 37 °C, added to liposomes and acidified to pH 5.1. After 1 min, the sample was neutralized to pH 8.0. The sample was incubated with 0.25% β-ME for 30 min at 37 °C, trypsinized for 1 h and subjected to SDS-PAGE analysis. CHK-152 concentration at incubation: ++, 20 nM CHK-152 in ratio of 335 to virions. Representative image out of 3 experiments is shown.

Radiolabeled CHIKV pre-incubated with 10 nM CHK-152 was added to liposomes after which the mixture was acidified to pH 5.1 for 1 min and back-neutralized to pH 8.0. A sucrose density column was formed from a layer of 60% (w/v) sucrose, then the sample mixed with 50% sucrose, and on top of that 20% and 5% layers. Upon ultracentrifugation, liposome-bound virus particles are at the 5– 20%-layer interface, whereas unbound particles remain within the 50% sucrose layer. The radioactivity counts were determined, providing a measure of virus co-floating with, and therefore bound to, the liposomes. In the absence of antibodies, on average 55% binding was observed that was set to 100%. Comparable virus-liposomes binding was observed in the presence of an isotype antibody. Importantly, however, virus-liposome binding was completely abolished in presence of CHK-152 antibodies. This observation suggests that CHK-152 prevents stable interaction of E1 to liposomes and as a consequence no membrane fusion is observed.

To investigate if CHK-152 indeed blocks the low-pH induced conformational changes that are required for membrane fusion, we assessed the formation of a trypsin-resistant form of E1 under low-pH conditions (Figure 2b). It is known that the E1 homotrimer of alphaviruses that is formed upon low-pH treatment is resistant to trypsin digestion (Kielian, Helenius, 1985). The trypsin-resistant E1-trimer dissociates into monomers when boiled in SDS sample buffer and can be detected with SDS-PAGE analysis. CHK-152-opsonized, radiolabeled CHIKV was incubated with liposomes at pH 5.1 as described for the liposome-binding assay (also see Methods). After back-neutralization to pH 8.0, the acidified liposome-CHIKV mixture was incubated with the reducing agent β-mercaptoethanol for 30 min in order to make the proteins more accessible to trypsin cleavage. The sample was then subjected to trypsin digestion. As expected, in the absence of CHK-152, a clear trypsin-resistant E1-band is seen. In presence of 20 nM CHK-152, however, the formation of the trypsin-resistant form of E1 was markedly reduced. Collectively, these observations suggest that high concentrations of CHK-152 either freeze the particle in the original state or interfere with an early step in the membrane fusion process i.e. at a step prior to stable interaction of E1 with the target membrane.

### The single-particle assay

We established that CHK-152 blocks efficient membrane interaction both at neutral and low pH at high antibody concentrations. At lower antibody concentrations, however, CHIKV was able to bind to planar bilayers (Figure 1a) and we aimed to elucidate if at these conditions CHK-152 is able to directly interfere with membrane fusion, and if so, to determine the stoichiometry of CHK-152 mediated neutralization of membrane fusion. To this end, we employed a single-particle assay with fluorescently tagged CHK-152, enabling counting of the number of CHK-152 bound to the individual viral particles. The single-particle assay relies on a controlled in vitro environment that enables synchronized acidification to initiate fusion and uses fluorescent tags to correlate the rate and extent of fusion to antibody binding.

The essentials of the single-particle assay are illustrated in Figure 3. The features were similar to those described before (van Duijl-Richter et al., 2015, Otterstrom et al., 2014). The basis is an in vitro flow cell system that allows rapid acidification of virions that are pre-docked onto a planar lipid bilayer (Figure 3a), monitoring at the same time for every particle the occurrence of hemifusion and the number of antibodies present. As described above, a planar lipid bilayer was formed on a hydrophilic coverslip in a flow cell. A biotinylated lipid provided an anchor for fluorescein-labeled streptavidin to report on the change in local pH. CHIKV particles were membrane-labeled with the lipophilic dye R18, incubated at 37 °C with or without antibody, and flown into the flow cell to dock nonspecifically to the bilayer. After acidification, hemifusion was observed as the escape of R18 from the viral membrane into the target bilayer (Figure 3b), and the time from pH drop to hemifusion was determined.

**Figure 3.**
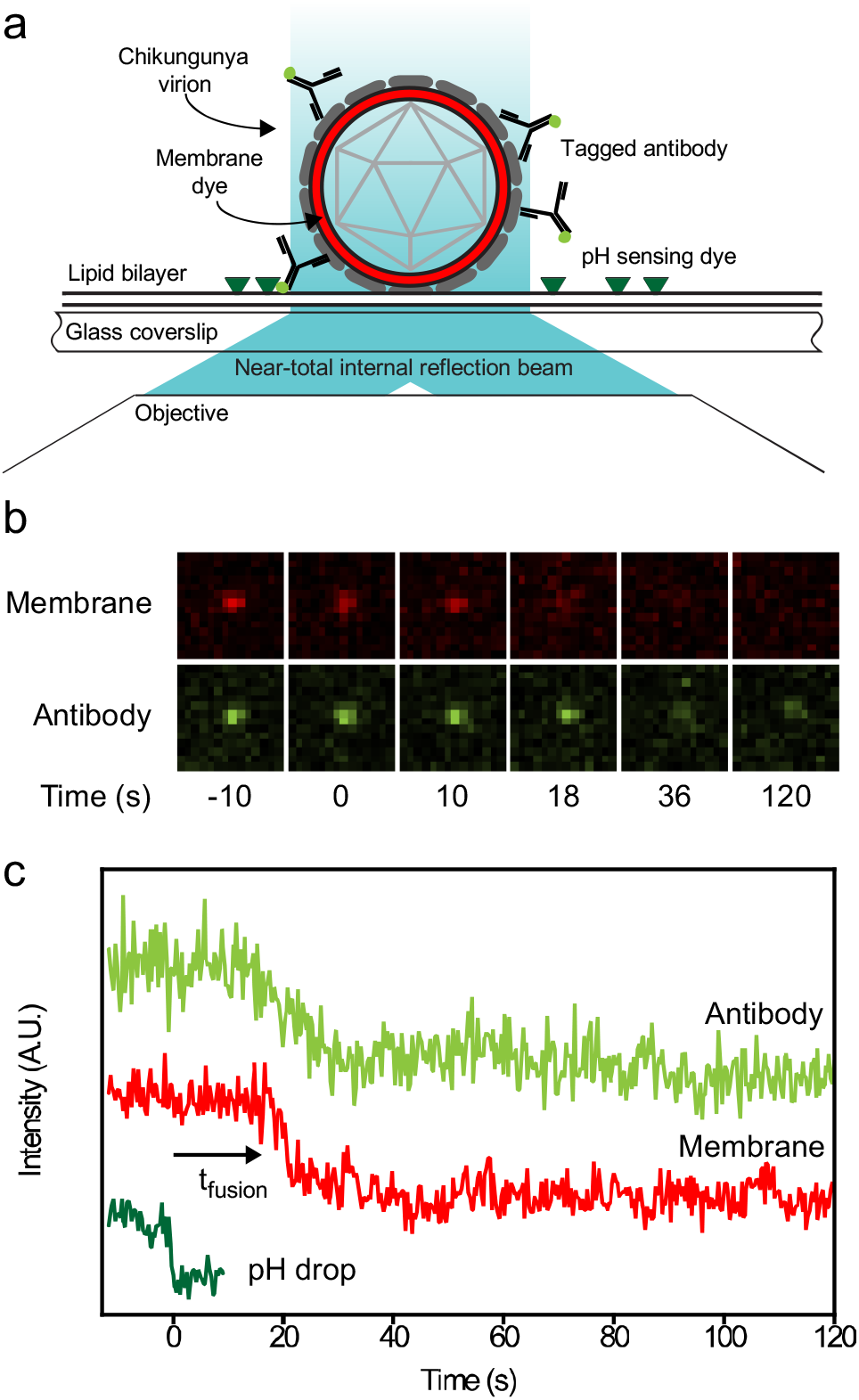
Single-particle assay. (a) In a flow channel, a lipid bilayer was formed on a cover glass. Viruses were labeled with lipophilic dye R18 and docked nonspecifically. A pH-sensitive dye attached to the membrane reported on pH change in the channel. Antibodies were detected and counted through a fluorescent tag. Fluorescence was excited by laser beams leaving the coverslip at a small angle. Fluorescence was split and projected onto different halves of a camera, allowing colocalization of the viral membrane and antibody spots. (b) Examples of observed fluorescence (membrane and antibody) of the same virus particle. Hemifusion can be seen around 16 s after acidification as escape of the membrane dye into the target bilayer. Loss of antibody intensity is also observed. (c) Intensity information collected from the virus particle in panel b. Top trace shows the loss of antibodies over time after acidification. Middle trace shows the membrane intensity signal. The lower trace shows the disappearance of fluorescence of the fluorescein pH probe, defining the start of the experiment. The time to hemifusion, defined as the onset of signal dissipation, is indicated as *t*fusion.

### CHK-152 blocks and slows down fusion of pre-docked virions in a pH-dependent manner

To correlate the effect of CHK-152 to different fusion conditions, we determined the fusion extent and time to fusion at pH 6.2, 6.1, 5.1 and 4.7. The latter two pH points lie in the optimal regime of fusion, and the first two around the threshold of fusion activation (see Figure 4– Figure supplement 1 and (van Duijl-Richter et al., 2015)). Measurements at pH 6.2 and 6.1 are in the pH range of early endosomes from which CHIKV particles have been described to fuse (Hoornweg et al., 2016). We studied fusion at room temperature; the rate of fusion scaled in an Arrhenius-like fashion over the range 37 °C to room temperature as determined with the liposomal fusion assay described above (Figure 4– Figure supplement 2). The extent of fusion, the fraction of the particle population that undergoes hemifusion within 2 min after acidification, is shown in Figure 4a.

**Figure 4.**
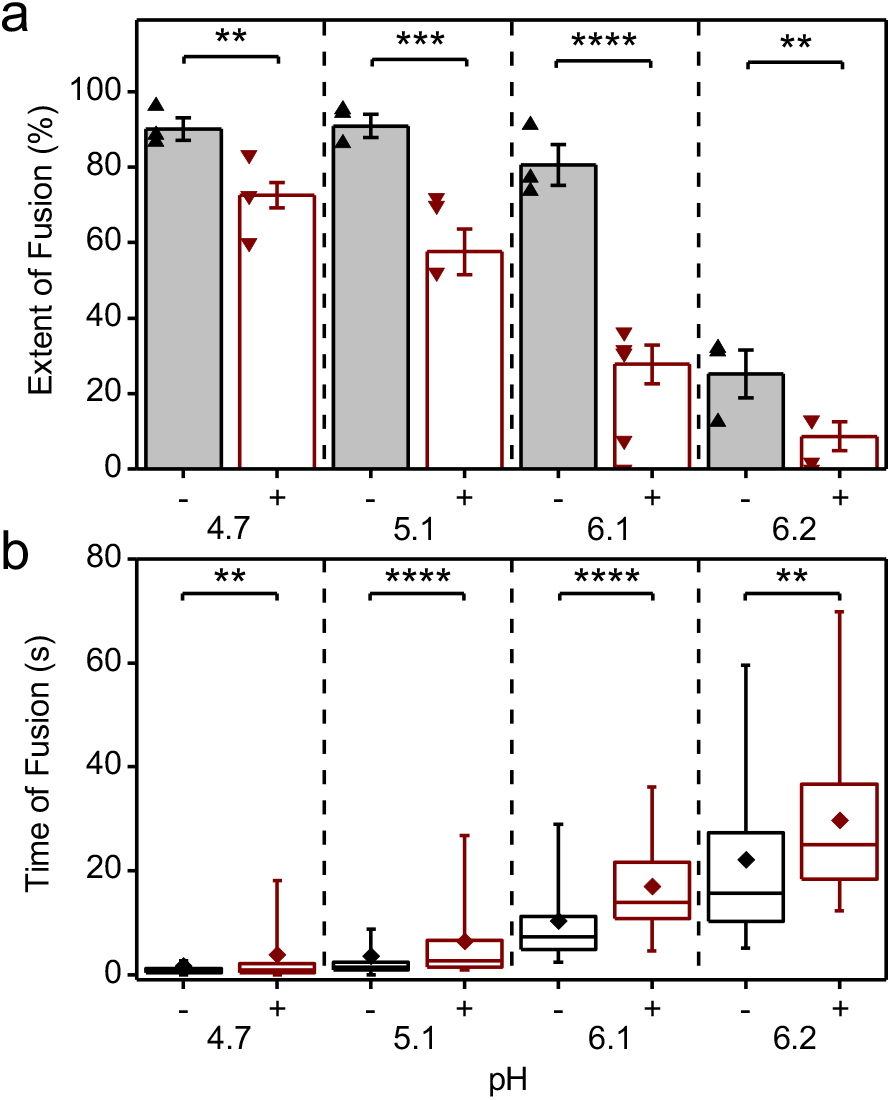
Inhibition and slow-down of CHIKV fusion by CHK-152, in a pH-dependent manner. (a) Virions pre-docked to the planar bilayer were acidified to the pH-point indicated below the x-axis, either with (+) or without (-) pre-incubation with CHK-152. The extent of fusion, the fraction of the population undergoing fusion, is shown. Mean±sem shown together with single experiments (triangles): black/-, without CHK-152, red/+, with pre-incubation of 0.63 nM CHK-152, resulting in 52±3 CHK-152 bound (see text). Significances determined by weighted t-test. (b) Time of hemifusion of single particles with the same color coding of conditions as panel a. Means, diamonds; box plots, 5%-Q1-median-Q3-95% intervals. Significance of difference of medians determined by Wilcoxon rank-sum test. Obtained p-values (Figure 4– Table supplement 1) **: p<0.01, ***: p<0.001, ****: p<0.0001. pH 6.1 and 6.2 lay at the threshold of fusion (Figure 4– Figure supplement 1). Fusion was studied at room temperature as the rate of fusion scaled in an Arrhenius-like fashion over the range 37 °C to room temperature as determined with the liposomal fusion assay (Figure 4– Figure supplement 2). Figure 4– Figure supplement 3 details the CHK-152 numbers bound and shows no correlation between the starting number of CHK-152 and the fate of fusion. There was some antibody-induced virion aggregation and therefore virions with high antibody counts were filtered out (Figure 4– Figure supplement 4 and Methods). Figure 4– Movie supplements 1 through 8 show representative timelapses of each condition.

Fusion was highly efficient, with experiments showing up to 96% extent of fusion. As previously observed for the S27 strain (van Duijl-Richter et al., 2015), the LR2006-OPY1 strain exhibited a sharp pH threshold between pH 6.2 and 6.1, with the extent of fusion reduced by half over a pH difference of 0.1. The time to hemifusion of single particles is shown in Figure 4b and shows that the time to fusion is longer with higher pH.

CHK-152 was labeled with AlexaFluor488 to enable quantification of the copy number bound to single virions. To this end, both the intensity of single, tagged CHK-152 and the unlabeled fraction of antibody were determined (Methods). Because CHK-152 incubation induced some amount of virion aggregation, we analyzed 75% of the virus particles, those with the lowest antibody counts (more details in Methods). CHIKV was incubated with 0.63 nM of tagged CHK-152 for 15 min at 37 °C to allow binding to occur. This concentration resulted in an average of 52±3 antibodies bound per virion with minor preparational variation per pH condition (Figure 4– Figure supplement 3a). This number corresponds to 22–43% of the 240 epitopes bound depending on the valency of CHK-152 binding (see Discussion). Under all conditions, this number of bound CHK-152 reduced the total extent of fusion (Figure 4a), indicating that CHK-152 directly blocks fusion at concentrations leading to sub-maximum epitope occupancy. The largest relative inhibition was observed at the threshold pH of 6.1 and 6.2. In addition to a reduction in extent, fusion was slowed down significantly under all pH conditions (as tested on the medians, Figure 4b). There was no consistent correlation between fusion of particles and starting antibody count (Figure 4– Figure supplement 3b). This observation may indicate that only a small number of the CHK-152 bound determine the fate of fusion, a number small enough that it does not contribute a detectable correlation.

### CHK-152 dissociates from viral particles at low pH

We observed that at pH 4.7 and 5.1 the fusion inhibition was reduced compared to the pH 6.1 and 6.2 conditions even though the initial binding levels of CHK-152 were similar (Figure 4– Figure supplement 3a). Hence, we decided to check the amount of CHK-152 bound to the virus particles over time. Figure 5a shows observed spots from single virions bound with fluorescently tagged CHK-152. After 2 min at pH 4.7, almost all fluorescence had disappeared from spots of non-fusing virions, indicating CHK-152 dissociation. In contrast, at pH 6.1 only marginal reduction of fluorescence was observed.

The average bound number of CHK-152 over time was determined for fusing and non-fusing Adj. R-particles separately (Figure 5b and Figure 5– Figure supplement 3). Time *t* = 0 was defined by the loss of fluorescence of the pH-sensitive fluorescein, and signals showed an initial increase towards *t* = 0 due to the rolling and arrest of virions under the force of the inflowing low-pH buffer. Both fusing and non-fusing virions displayed CHK-152 dissociation at pH 5.1 and 4.7. Because fusing particles additionally lost CHK-152 after fusion due to diffusion (Figure 5a, top images) we decided to take the number of CHK-152 bound to the non-fusing particles (Figure 5b) as a proxy for the dissociation behavior of the whole population, as this indicates purely dissociation into solution.

**Figure 5.**
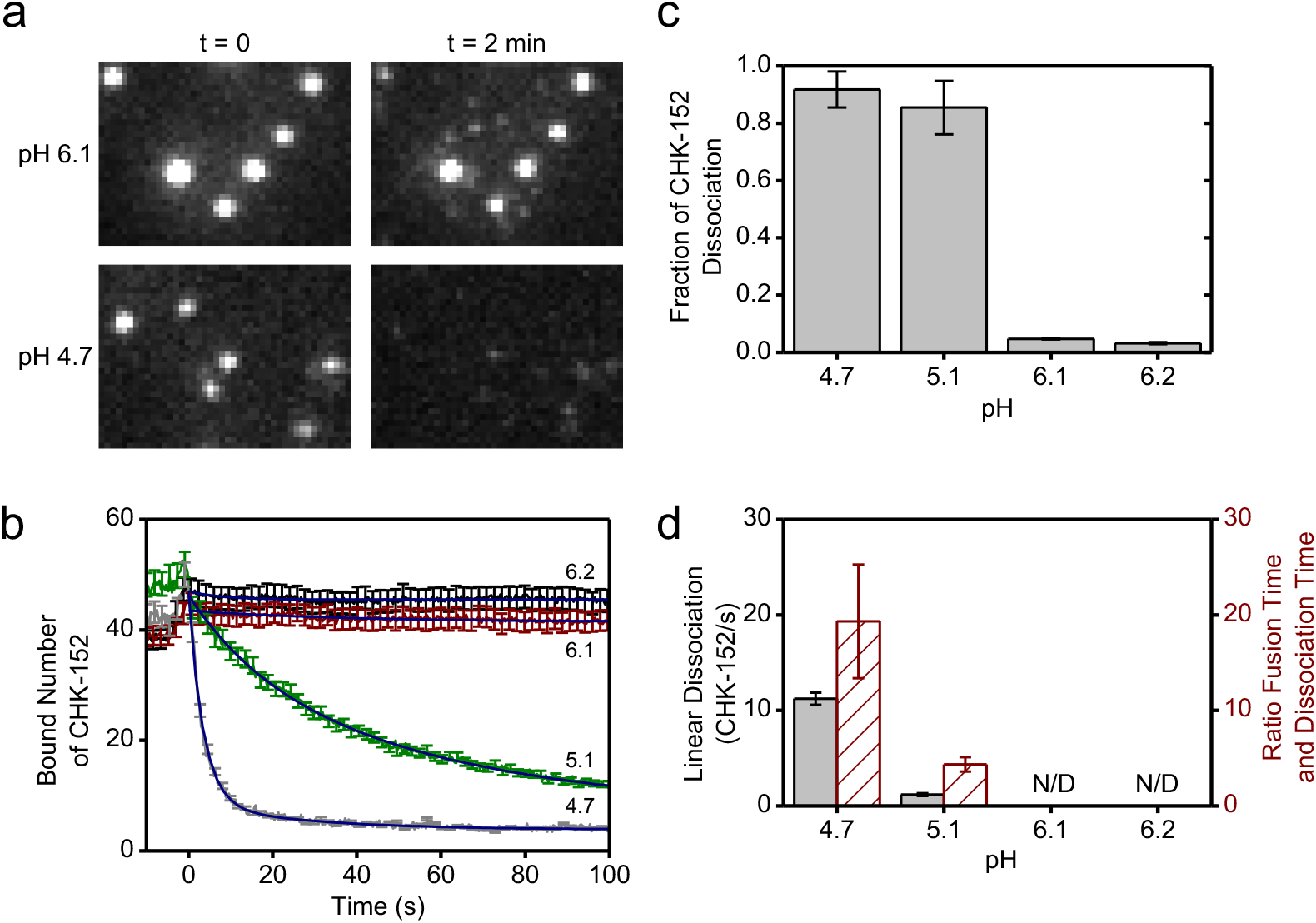
CHK-152 dissociation at low pH. (a) Fluorescent spots of CHK-152 bound to virions are shown from a region of a movie slice for pH 6.1 and 4.7, and for *t* = 0 and *t* = 2 min. At pH 4.7, losEsqouafti CHK-152 from the virions was observed after 2 min. Image heights correspond to 8.5 μm. (b) The average of bound CHK-152 of non-fusing virions is shown over time. Increase of signal towards *t* ^R^=^e^0^duc^ was due to rolling and arrest of virus particles. One out of every five error bars shown to reduce visual clutter. Blue lines show exponential fits (see text). (c) The final fraction of antibody remaining for each pH point was determined from the fits in panel b. (d) The linear rate of dissociation at *t* = 0 determined from the fits in panel b is shown per pH point in black (left y-axis). Red bars (right y-axis) show the ratio of the mean fusion time without antibody (see Figure 4) to the dissociation time (_B_th_ou_e_nd_ inverse of the linear dissociation rate), at the pH points indicated. All error bars, sem. N/D: not Model detectable. Confirmation of CHK-152 labeling in Figure 5– Figure supplement 1. Single CHK-152 intensity determination in Figure 5– Figure supplement 2 and Methods. The average of bound CH^Eq^K^u^-^ati^ 152 of fusing virions is shown in Figure 5– Figure supplement 3.

As the fusion yields were slightly different for pH 5.1 and 4.7, we determined the properties of CHK-152 dissociation for both pH points. The curves showing the number of bound CHK-152 over time were fit with single-exponential (pH 6.2 and 6.1) and double exponential (pH 5.1 and 4.7) decay functions to extract the fraction of CHK-152 that ultimately dissociate (Figure 5c). Only marginal loss of antibody was observed at pH 6.2 and 6.1, whereas more than 80% of antibodies dissociated at pH 5.1 and 4.7. From the fits, the linear rate of dissociation at *t* = 0 was determined for pH 5.1 and 4.7 (Figure 5d, red), showing that pH 4.7 features an about 10-fold faster initial dissociation rate. Importantly, the ratio of the rates of fusion and rates of dissociation differed (Figure 5d, green): at pH 4.7, CHK-152 dissociation is about 10-fold faster than at pH 5.1, while the mean fusion time is only about 2-fold faster. The rate of dissociation may therefore explain the differences in extent of fusion at pH 5.1 and 4.7. We postulate fusion would be blocked with the starting CHK-152 counts (like at pH 6.1 and 6.2). However, due to sufficiently fast dissociation, compared to the timescale of the events leading to fusion and E1 protein inactivation, some virions become fusogenic again. Dissociation happens more quickly at pH 4.7 than at 5.1 relative to the events that lead to membrane fusion, thereby leading to a higher fusion extent. We therefore numerically modeled the process leading to the observed fusion extents, taking the CHK-152 stoichiometry and dissociation into account.

### Antibody stoichiometry indicates high cooperativity at the level of E1/E2 spikes

Binding of CHK-152 blocked and slowed down fusion. However, most epitopes were not bound with CHK-152, and at pH 4.7 CHK-152 dissociated very fast. To explain how small numbers of antibody can inhibit fusion, we devised a numerical model of fusion in which a single CHK-152 bound to an E2 surface epitope prevents the whole spike from participating in fusion. This model bears semblance to earlier work by us and others on influenza fusion inhibition (Otterstrom et al., 2014, Ivanovic, Harrison, 2015). Also, dissociation of all CHK-152 bound to the spike would restore that spike’s fusogenicity, provided the dissociation happened rapidly enough compared to the fusion timescale. The fusion extent was then numerically evaluated by assessing the availability of a sufficient number of unbound spikes that are in contact with the target membrane. Comparison of the results of this model to the observed stoichiometries and dissociation properties can then inform us on the cooperativity of CHIKV fusion at the spike level.

The key parameters in the model were the total number of spikes associated with the target membrane and the number of spikes that need to cooperatively act to mediate fusion. We considered different sizes for the contact patch in interaction with the target membrane, containing *M* proteins (Figure 6a). A spike was considered not to participate in mediating fusion if one or more of the three spike epitopes were bound by antibody (Figure 6b). Fusion could only be attained if a virus particle had a number *N*_H_ of unbound spikes within any 5- or 6-ring in its contact patch. Here, *N*_H_ = 1 signifies fusion mediated by a single E1 trimer formed from an unbound spike, and for higher *N*_H_ fusion results from multiple unbound spikes in a ring on the viral surface (illustrated in Figure 6c). The positions of unbound spikes within the ring did not matter, as long as any ring in the contact patch contained *N*_H_ unbound spikes.

**Figure 6.**
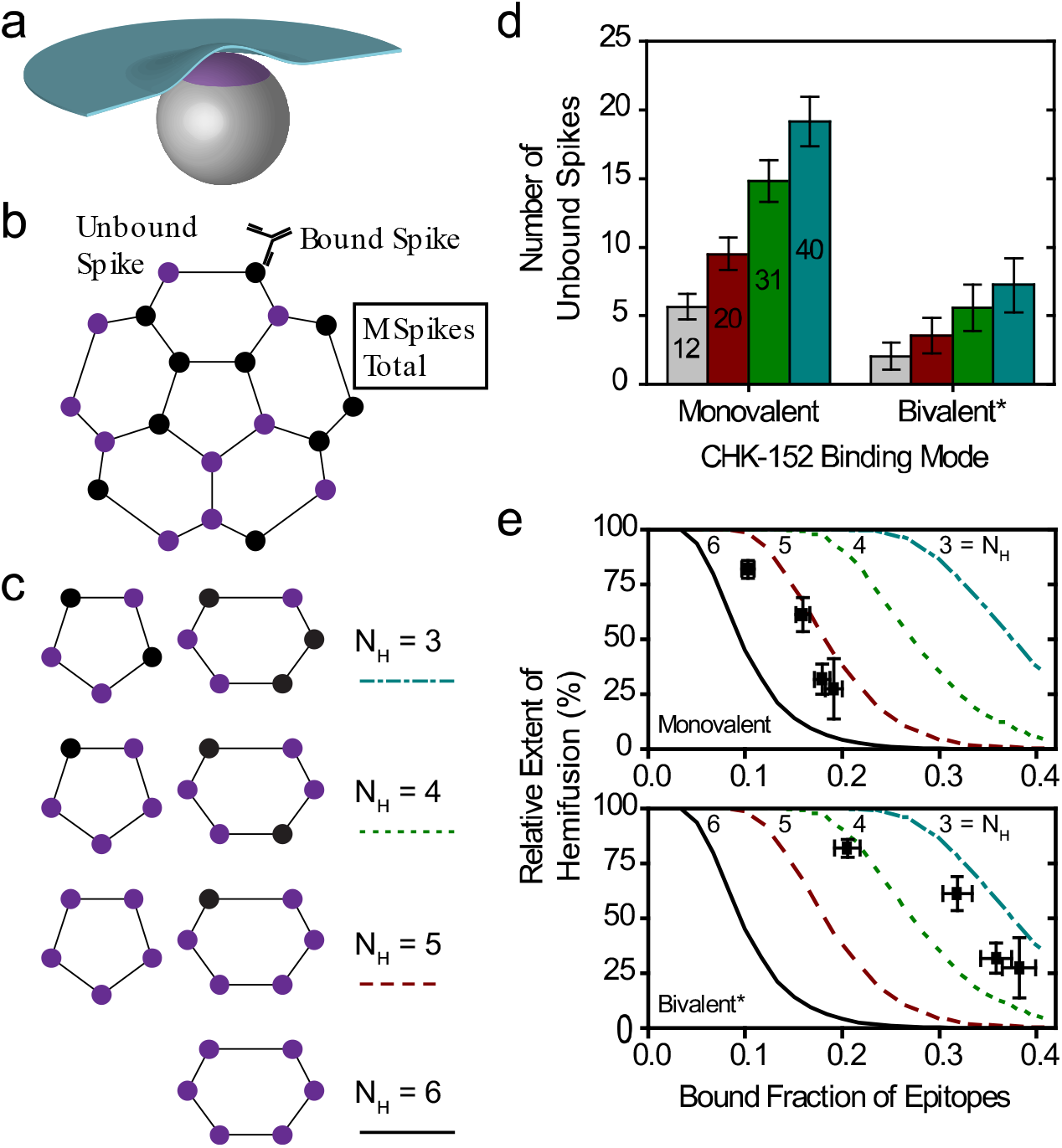
Cooperative model of CHIKV fusion at the level of spikes. (a) A virion (grey) docked to the planar membrane (blue) is shown. The region in contact with the target membrane is shown in purple: the contact patch. (b) The contact patch consists of *M* spikes, example of *M* = 20 shown. More details of the patch size are in Figure 6– Figure supplement 1. Unbound spikes (purple) are considered to mediate fusion whereas spikes bound with one or more CHK-152 are considered not to (black). (c) Cooperative fusion was modeled by the availability of a minimum number of unbound spikes, *N*_H_, in any of the 5- and 6-rings on the viral surface. The unbound positions can be anywhere in the ring; examples for different *N*_H_ are shown. (d) For 10 000 virions 52±3 CHK-152 were randomly bound per virion. Both the contact patch was varied (from 12 to 40) and the CHK-152 binding mode. The mean±SE of the number of unbound spikes is shown. Bivalent* binding was modeled as binding by 104±6 monovalent Fabs. (e) For 10 000 virions CHK-152 was randomly bound as in panel d and the relative extent of fusion was determined as the fraction of virions having available *N*_H_ free spikes in a ring as defined in panel c. The extents of fusion from the simulations are shown as lines versus the fraction of CHK-152-bound epitopes on the viral surface. Line legends are as shown in panel c: *N*_H_ = 3,4,5,6 are indicated by dash-dotted, dotted, dashed and a solid line respectively. The experimental extent of fusion was determined relative to the no antibody control (Figure 6– Figure supplement 2) and is plotted versus the time-averaged fraction of bound epitopes (black squares, mean±sem). This time-average takes into account CHK-152 dissociation (see text and Figure 6– Figure supplement 3). For panel e, different patch sizes and their influence on the best fit parameters is shown in Figure 6– Figure supplements 4 and 5.

We considered the two extreme cases of the CHK-152 binding mode with its two Fab domains: pure monovalent and pure bivalent binding. With a number of 52±3 antibodies bound over the 240 epitopes (in 80 spikes), the probability of a spike to be unbound is: *p*_unboundSpike_ = (1-52/240)^3^ = 0.48±0.03 for monovalent binding, or *p*unboundSpike = (1-104/240)^3^ = 0.18±0.03 for bivalent* binding. We write bivalent* binding, as this was estimated as binding of double the amount of monovalent Fabs. This is an unattainable maximum epitope occupancy, since bivalent antibodies can only bind neighboring epitopes and additionally will experience steric hindrance. Considering the probabilities calculated above, any contact patch of size *M* > 5, corresponding to greater than 6.25% of the virion surface, on average has more than 1 unbound spike in contact with the target membrane.

For a virion of 65 nm in diameter we estimate the contact patch to be 20 spikes, or 25% of the viral surface by looking at the range that the 13-nm-long E1 (Voss et al., 2010) may reach to a planar target membrane (Figure 6– Figure supplement 1a). Earlier work has similarly estimated the contact patch area of spherical, 50-nm diameter influenza viruses at 25% of the outer surface (Ivanovic et al., 2013). Here, the contact patch could be larger if inserting E1 were to pull the target membrane around the virion like a coat, or could be smaller due to steric hindrance of antibodies. In the biological context, the contact area with the inversely curved endosome may increase the contact patch. Therefore, we consider different sizes of *M* from 12 (about one eighth) to 40 (one half of a virion) as shown in Figure 6– Figure supplement 1b, which appear to be reasonable limits for the minimum and maximum contact patch size respectively. Then, we counted the number of unbound spikes in numerical simulations of the fusion. All tested patch sizes were determined to have multiple unbound spikes available on average (Figure 6d), in line with what we calculated above. We therefore considered a cooperative fusion mechanism.

First, we scaled the data to enable comparison with the numerical model. The extents of fusion in the presence of CHK-152 were calculated relative to the no-antibody condition, thereby correcting the extents for non-fusogenic virions and for the effect of pH on the total extent (Figure 6– Figure supplement 2). To correct for the dissociation of CHK-152 over time, we then calculated the effective number of CHK-152 bound to the virus particles during the time they fuse. We calculated this effective number over the timescale of fusion, by averaging the number of CHK-152 bound to non-fusing virions over the population, and subsequently averaging over time weighted by the number of particles that have not yet fused (see Figure 6– Figure supplement 3). It is therefore an estimate of the average number of CHK-152 a fusing virion had bound during the time to fusion. The result is shown in Figure 6e (squares): the observed relative extents of fusion versus the estimated effective epitope occupancies in the cases of monovalent and bivalent* binding.

Finally, we ran numerical simulations for 10 000 virions determining at each epitope occupancy what fraction of the virions had a ring containing *N*_H_ unbound spikes, defining the extent of fusion. The result is shown in Figure 6e as lines, for *M* = 20. We see that the data best matches fusion mediated by 3–5 unbound spikes in a ring (indicated by a red dashed and cyan dash-dotted line respectively), depending on CHK-152 binding valency. The cooperativity was largely determined by the valency of CHK-152 binding; the actual contact patch simulated was of minor effect (Figure 6– Figure supplement 4 and Figure 6– Figure supplement 5).

## Discussion

Here, we reported on the mechanism of action of antibody CHK-152. We determined that it shields the virions at high concentrations of binding, preventing membrane interaction under neutral-pH as well as low-pH conditions. Using a single-particle fluorescence assay and a sub-stoichiometric ratio of CHK-152 binding, virions were pre-docked to a membrane. This approach allowed us to determine that CHK-152 also plays a role in directly blocking the fusion step. In this assay, CHK-152 was observed to dissociate at low pH, whereas it remained bound at mildly acidic pH. We devised a numerical model of CHIKV fusion with only E1 from unbound spikes able to trimerize and mediate fusion, and in which fusion is achieved by insertion of a minimal number of E1 trimers within a ring of neighboring spikes. Correcting for CHK-152 dissociation, the CHK-152 stoichiometries of binding were not consistent with fusion by single E1 trimers, but rather with fusion mediated by three to five trimers.

In addition to CHK-152 effectively preventing viral docking to membranes at neutral pH, it appears to directly block low-pH fusion by interfering with stable attachment of the virus to the target membrane. Our data and previous work indicate that prevention of virus attachment to the cell, possibly by sterically hindering receptor or membrane interaction, is an important mechanism in its neutralizing efficiency (Pal et al., 2013). We demonstrated that CHK-152 also directly inhibits fusion for pre-docked virions, at sub-saturated occupancy of binding. This enhances its potential as an antiviral by the multiplicative effect of binding reduction and fusion inhibition. It has been shown before that the CHK-152 Fab binds residues in the E2 A domain and the β-ribbon. The latter lies in the acid-sensitive region that becomes disordered at low pH, facilitating exposure of the E1 fusion loop (Voss et al., 2010, Li et al., 2010, Sun et al., 2013). As we find that CHK-152 prevents the formation of a trypsin-resistant form of E1, and inhibits stable association of E1 with target membranes, it seems plausible that CHK-152 inhibits E1 membrane insertion by blocking E1-E2 heterodimer dissociation. However, it could also lock the E2 proteins in place allowing E1 membrane insertion but preventing trimerization, as observed in studies at threshold pH of 6.4 for Sindbis virus (Cao, Zhang, 2013). Interestingly, the acid-sensitive region and A and B domains appeared more often as binding targets for antibodies (Fox et al., 2015, Selvarajah et al., 2013, Smith et al., 2015). The epitope of neutralization lies within one single E2 molecule, in contrast with other, E2-crosslinking antibodies isolated for alpha- and flaviviruses (Jin et al., 2015, Fox et al., 2015, Kaufmann et al., 2010), so ‘locking’ the virion would require CHK-152 bivalent binding.

We observed CHK-152 dissociation at pH 5.1 and 4.7. In the in vitro conditions of our experiment, all unbound CHK-152 had been washed away so that CHK-152 dissociating after acidification effectively disappeared. This is in contrast with the liposomal fusion conditions (Pal et al., 2013) and an in-vivo situation, where CHK-152 might rebind from solution. Also, at the probed stoichiometry of binding in the single-particle assay, dissociation of just a couple of CHK-152 may restore virion fusogenicity. This would not be the case for higher concentrations of antibody incubation. Dissociation was marginal at pH 6.1 and 6.2, the pH of the early endosome through which CHIKV enters cells (Hoornweg et al., 2016), and the extent of fusion was strongly reduced at these pH points. Also, the CHIKV strains so far have a sharp pH threshold and appear to be liable to acid-induced inactivation (van Duijl-Richter et al., 2015). In all, CHK-152 dissociation may not need to compromise its neutralization effectiveness in vivo even at sub-stoichiometric binding levels.

We found that the relative rate of CHK-152 dissociation determined the final extent of fusion for pH 4.7 and 5.1. However, at both pH points nearly all CHK-152 dissociated if given enough time. Together, this indicates that there is a “window of opportunity” during which the spikes must become unbound in order to still be able to mediate fusion again. Such a window of opportunity may arise for example due to inactivation of E1 proteins at low pH, as observed without the presence of target membranes (van Duijl-Richter et al., 2015). Even though the window of opportunity is an underlying, necessary assumption of our model, we did not explicitly model it as we just considered the average presence of CHK-152 for particles during the time they take to fuse.

Two different mechanisms of CHK-152 dissociation could be involved. In the first, the CHK-152 lose affinity due to protonation changes in the epitope or paratope. This may involve an antibody-induced shift of the p*K*_a_ of protonatable residues on the protein, as suggested in Zeng et al (Zeng, Mukhopadhyay & Brooks, 2015). In the second, we see an analogue to how the influenza hemagglutinin has been modeled to overcome the kinetic barrier to rearrange to the post-fusion state by protonation (Zhang, Dudko, 2015). Here, the CHK-152 would raise the kinetic barrier for E2-E1 heterodimer dissociation. However, this increased barrier to conformationally rearrange is then overcome at sufficiently low pH, shedding the antibody. Identifying the dissociation mechanism is beyond this work as both described changes in CHK-152 and viral protein are proton-triggered. However, it appears important to determine if this mechanism is common in antibody-mediated neutralization of class II viruses, if it allows decreased-pH-threshold escape mutants to arise and if this could be avoided or exploited in rational antiviral design.

Employing the fusion-inhibiting capacity of CHK-152, we found CHIKV fusion to be cooperative by determining the stoichiometry of binding of CHK-152 and numerically simulating the resulting availability of CHK-152-free spikes on the virion surface. Fusion ensued when a sufficient number of unbound spikes were available to trimerize and together overcome the membrane fusion barriers. In this scenario, the E1 trimer fusion loops could associate to facilitate dimpling of both membranes, as detected before for the E1 ectodomain (Gibbons et al., 2004, Gibbons et al., 2003). The proposed mechanism is analogous to that developed for influenza viral fusion, where multiple protein trimers need to mediate fusion and the network of potentially cooperating trimers is disrupted by inhibitor binding (Otterstrom et al., 2014, Ivanovic, Harrison, 2015). Interestingly, in those studies, binding of an estimated quarter of epitopes resulted in significant fusion inhibition, similar to the occupancy probed here.

The combination of data and numerical model allowed to determine that CHIKV fusion is cooperative, but some uncertainties remain. To develop a more complete understanding of CHIKV fusion, it is necessary to probe a large range of inhibitor binding concentrations and obtain sufficient statistics to allow inference on the individual protein events to membrane fusion (for instance, the steps of heterodimer dissociation and E1 membrane insertion). The distribution of fusion times then allows inference on the underlying rate-determining steps (Ivanovic, Harrison, 2015, Chao et al., 2014, Kim et al., 2017). Here, we were limited to sub-stoichiometric levels of binding as CHK-152 prevented nonspecific membrane docking at high binding levels, and the statistics were too limited to determine the fusion time distributions. The actual number of E1 trimers involved in fusion depended for the most part on the valency of the CHK-152 and less on the size of the contact patch. We point out two additional factors why CHIKV fusion is more cooperative than we could probe. First, the CHK-152 inhibited nonspecific docking, and virions may therefore have preferentially bound with a relatively sparsely CHK-152-covered section of the viral surface. The epitope occupancy in the contact patch is then relatively lower than on the rest of the particle, which implies a more cooperative fusion mechanism. Second, we see no reason a priori why E1 from different spikes would be prevented from forming a trimer together. Compared to our model, this would further increase the number of E1 trimers that could form in the contact patch, thereby also implying a more cooperative mechanism.

Because of the reasons stated above, future studies should uncouple the binding- and fusion-inhibiting action of inhibitors by artificially coupling viruses to the membrane surface. Furthermore, using monovalent-binding Fab fragments eases interpretation of the data, and may reduce steric effects. Our results on alphavirus fusion fit in with a universal context found so far across all three classes of enveloped viruses, where fusion is mediated by multiple protein trimers in a close neighborhood (Ivanovic et al., 2013, Chao et al., 2014, Kim et al., 2017). Taken together, our data identifies important parameters to consider in the rational development of CHIKV antivirals.

## Materials and Methods

CHIKV strain LR2006-OPY1 was a kind gift by Dr Andres Merits. Antibody CHK-152 was a kind gift from Dr Michael Diamond. All assays were performed at 37 °C, except the single-particle assay which was performed at room temperature (around 22 °C). The corresponding change in the rate of fusion was determined in the liposomal fusion assay described below (Figure 4– Figure supplement 2). Throughout this Chapter we refer to (hemi)fusion as fusion, as the assays used do not distinguish content mixing from lipid mixing. Appendix contains details of hypothesis testing (Figure 4– table supplement 1) and fitting (Figure 5– table supplement 1).

Virus – radiolabeled. A confluent layer of BHK-21 cells was infected at an MOI of 10. The virus inoculum was removed after 2.5 h incubation and following a 1.5 h starvation, 200 μCi (7.4 MBq) [35S]-methionine/L-[35S] cysteine using EasyTag™ EXPRESS35S Protein Labeling Mix (PerkinElmer, Groningen, the Netherlands) was added to the medium. Supernatant was harvested 20 hpi (hours post-infection) and layered on top of a two-step sucrose gradient (20%/50% w/v in HNE) and centrifuged for 2 h at 154 000 x g at 4 °C in a SW41 rotor (Beckman Coulter, Woerden, the Netherlands) to clear from cell debris. Radioactive virus was collected at the 20%/50% sucrose interface and radioactivity was counted by liquid scintillation analysis. Fractions were pooled based on radioactivity counts. The infectivity of the virus preparation was determined by standard plaque assay on Vero-WHO cells and by qRT-PCR to determine the number of genome-containing particles, as described before (van Duijl-Richter et al., 2015).

Virus – fluorescently labeled, and inactivated. Virus stocks were prepared as described before (van Duijl-Richter et al., 2015). Briefly, CHIKV seed stocks were prepared by infection of Vero-WHO cells at a multiplicity of infection (MOI) of 0.01. Pyrene-labeled virus was produced in BHK-21 cells cultured beforehand in the presence of 15 μg/ml 1-pyrenehexadecanoic acid (Thermo Fisher Scientific, Waltham, MA, USA). The supernatant was harvested at 48 hpi, cleared from cell debris by low-speed centrifugation, purified by ultra-centrifugation and frozen in liquid nitrogen. Before freezing, the virus was UV-inactivated as the single-particle fusion assay was performed outside the BSL-3 facility (van Duijl-Richter et al., 2015). To produce octadecyl rhodamine B chloride (R18; Thermo Fisher Scientific)-labeled virions, 7.2×10^12^ particles of purified and inactivated CHIKV were diluted in PBS (10 mM phosphate, 140 mM NaCl, 0.2 mM EDTA) and 0.3 μL of 0.2 mM R18 dissolved in DMSO was added to a final concentration of 1 μM. Subsequently, the virus solution was kept on ice for 1 h. A gel-filtration column (PD-10 desalting column; GE Healthcare, Hoevelaken, the Netherlands) was used to separate the virus from unincorporated dye. The most concentrated fractions were combined and used in the experiment.

Liposomes. Liposomes were prepared as described before (van Duijl-Richter et al., 2015, Smit, Bittman & Wilschut, 1999). For the non-single-particle assays, the liposomes consisted of sphingomyelin from porcine brain, transphosphatidylated L-α-phosphatidylethanolamine (PE) from chicken egg, L-α-phosphatidylcholine (PC) and cholesterol from ovine wool. The lipids were mixed in a molar ratio of 1:1:1:1.5. The liposomes were prepared by freeze-thaw extrusion and extruded through a polycarbonate membrane with 200 nm pore. All lipids and the polycarbonate membrane were purchased from Avanti Polar Lipids (Alabaster, Alabama, USA). Lipids and the phospholipid-to-cholesterol-ratio were chosen to approximate the lipid composition within the endosomal compartment (Kolter, Sandhoff, 2010, van Meer, Voelker & Feigenson, 2008). For the single-particle assay, liposomes (200 nm) were also prepared by freeze-thaw extrusion. Liposomes consisted of 1:1:1:1.5:2×10-5 ratio of 1,2-dioleoyl-sn-glycero-3-phosphocholine (DOPC), 1,2-dioleoyl-sn-glycero-3-phosphoethanolamine (DOPE), porcine brain sphingomyelin (SPM), ovine wool cholesterol and 1,2-dioleoyl-sn-glycero-3-phosphoethanolamine-N-(biotinyl) (Biotin-PE).

Trypsin cleavage of CHIKV structural proteins at neutral pH. [35S]-methionine/L-[35S] cysteine labeled CHIKV was incubated for 10 min at 37 °C with CHK-152 in HNE in the appropriate ratio. In final volume for the tested conditions: 0.63 nM CHK-152 in approximate ratio of 13 to virions, and 10 nM CHK-152 in ratio of 210 to virions. Thereafter, liposomes were added at a final concentration of 200 μM at 37 °C in a total volume of 133 μL HNE buffer (5 mM HEPES, 150 mM NaCl, 0.1 mM EDTA) and kept for 60 s at pH 7.4.The mixture was digested with N-tosyl-L-phenylalanyl chloromethyl ketone (TPCK)-treated trypsin (Sigma-Aldrich, St. Louis, MO, USA) at a concentration of 200 μg/mL in the presence of 1% Triton X-100. After 1 h at 37 °C, the samples were subjected to SDS-PAGE analysis.

Trypsin cleavage of E1 homotrimer at low pH. [35S]-methionine/L-[35S] cysteine labeled CHIKV was prior opsonized with CHK-152: 20 nM CHK-152 in ratio of 335 to virions. This was then mixed with 200 μM liposomes at 37 °C in a total volume of 133 μL HNE buffer (5 mM HEPES, 150 mM NaCl, 0.1 mM EDTA). After 60 s of incubation, the pH was lowered to pH 5.1 by the addition of 7 μL of a pre-titrated buffer (0.1 MES, 0.2 M acetic acid, NaOH to achieve desired pH). After 60 s, the mixture was neutralized to pH 8.0 by the addition of 3 μL of pre-titrated NaOH solution. Samples were incubated in 0.25% β-mercaptoethanol (β-ME) for 30 min and subsequently digested with TPCK-treated trypsin (Sigma) at a concentration of 200 μg/mL in the presence of 1% Triton X-100. Samples were then subjected to SDS-PAGE analysis.

SDS-PAGE analysis. Samples were solubilized by 4x SDS sample buffer (Merck-Millipore, Darmstadt, Germany) and analyzed by SDS-PAGE on 10% Mini-PROTEAN^®^ TGX^™^ Precast Protein Gels (Biorad, Hercules, CA, USA). Gels were fixed in 1 M sodium salicylate for 30 min and dried. Viral protein bands were visualized in a Cyclone Plus Phosphor Imager (PerkinElmer) and radiographs were further analyzed using ImageQuant.

Liposomal binding assay. The influence of antibody binding of CHIKV on low-pH induced liposome-binding was assessed using a liposomal binding assay described before for SFV and SINV (Wahlberg et al., 1992, Smit, Bittman & Wilschut, 1999, Bron et al., 1993). Briefly, 0.75 μM viral phospholipid of [35S]-methionine/L-[35S]-cysteine labeled CHIKV particles was mixed with 200 μM liposomes in HNE buffer. The mixture was acidified by adding a pre-titrated amount of low pH buffer (0.1 M MES, 0.2 M acetic acid, NaOH to achieve desired pH). At 60 s after acidification, the mixture was neutralized to pH 8.0 by NaOH and placed on ice. 100 μL of this fusion reaction was added to 1.4 mL of 50% sucrose in HNE (w/v). A sucrose density gradient was prepared consisting of 60% sucrose in HNE, followed by 50% sucrose in HNE including the fusion mixture, 20% sucrose in HNE and 5% sucrose in HNE on top. Gradients were centrifuged in a SW55 Ti rotor (Beckman Coulter) for 2 h at 150 000 × g. The gradient was fractionated in ten parts and radioactivity in each fraction was determined by liquid scintillation analysis. The relative radioactivity in the top four fractions compared to total radioactivity in the gradient was taken as the measure for CHIKV that were bound to liposomes. For antibody-inhibition, [35S]-methionine/L-[35S]-cysteine labeled CHIKV was incubated for 10 min at 37 °C with 10 nM of CHK-152 in HNE before proceeding with a fusion measurement as described above.

Single-particle fusion – assay and microscopy. Experiments were performed at room temperature as reported before (van Duijl-Richter et al., 2015, Otterstrom et al., 2014). Glass microscope coverslips (24 mm x 50 mm, No. 1.5; Marienfeld brand, VWR, Amsterdam, the Netherlands) were cleaned using 30 min sonications in acetone and ethanol, followed by 10 min sonication with 1 M potassium hydroxide and finally 30 min cleaning in an oxygen plasma cleaner. The last step was performed on the day of measurement. Polydimethylsiloxane (PDMS) flow cells with a channel cross-section of 0.1 mm^2^ were prepared as before (Otterstrom et al., 2014). Imaging was performed with near-total internal reflection fluorescence microscopy (TIRF-M), using an inverted microscope (IX-71, Olympus, Leiderdorp, the Netherlands) and a high numerical aperture, oil-immersion objective (NA 1.45, 60×; Olympus). Liposomes were flushed into the flow cell and a planar lipid bilayer was allowed to form for >50 min. Virions were docked non-specifically to the lipid bilayer for 3 min at 50 μL/min. Fluorescein-labelled streptavidin (Thermo Fisher Scientific) was introduced into the flow cell at 0.2 μg/mL for 5 min at 10 μL/min, as a pH drop proxy. Then, PBS with 2 mM Trolox ((±)-6-Hydroxy-2,5,7,8-tetramethylchromane-2-carboxylic acid, Sigma-Aldrich) was flown in for 2 min at 100 μL/min to remove unbound virions and fluorescein. The presence of Trolox prevented laser-intensity dependent fusion inactivation, presumably by reducing oxidative damage from the fluorescent dye. The aqueous environment was acidified by flowing in citric acid buffer (10 mM, 140 mM NaCl, 0.2 mM EDTA) of pH 5.1 at 600 μL/min. The fluorophores were excited using 488 nm and 561 nm lasers (Sapphire, Coherent Inc., Santa Clara, CA, USA). Viral membrane fluorescence (red) and fluorescein pH drop fluorescence (green) were projected on different halves of an EM-CCD camera (C9100-13, Hamamatsu, Iwata-shi, Shizuoka-ken, Japan). Exposure time was 300 ms. Opsonization was performed for 15 min at 37 °C with appropriate concentration of antibody and 10x diluted labeled virus, in final volume.

Antibody labeling and characterization. CHK-152 was labeled with AlexaFluor488 TFP-ester (Thermo Fisher Scientific) per manufacturer’s guidance. UV-VIS spectroscopy indicated a labeling ratio of 1.5 dye/CHK-152. Tandem MALDI mass spectrometry was consistent with this (Figure 5– Figure supplement 1). MALDI was performed in 150 mM ammonium acetate, after dialysis. From the labeling ratio we estimated the fraction of unlabeled (i.e., not visualized) CHK-152 at 0.22, by assuming a Poissonian labeling distribution. To determine single CHK-152 intensity, labeled CHK-152 was flown in at roughly picomolar concentration into a clean flowcell as described above. Imaging conditions and buffers were the same as for virions (i.e. pH 5.1, unless noted otherwise). Single CHK-152 intensity was determined in a 7×7-pixel region, to be 36±2 A.U. per CHK-152 (Figure 5– Figure supplement 2a), corrected for background and laser intensity. Antibody fluorescence intensity was independent of pH (Figure 5– Figure supplement 2b).

Single-particle fusion – analysis. Home-written software in MATLAB was used to extract the fluorescence signals, essentially as described before (van Duijl-Richter et al., 2015, Otterstrom et al., 2014). In brief, the fluorescein pH-drop signal was integrated over the entire field of view and the t=0 of the experiment defined as the time point where only 8% of a fitted sigmoidal function remained. Particles fusion events and times were manually detected by inspecting the virion R18 intensity traces together with the movie. CHK-152 fluorescence traces were extracted in a 7×7-pixel region, corrected for background, laser intensity and laser illumination profile, and divided by the intensity per CHK-152 and dark fraction as determined above, to yield the number of CHK-152 bound. As we detected virion aggregation, presumably by antibody crosslinking, in both an increased R18 intensity distribution and a bimodal CHK-152 distribution (Figure 4– Figure supplement 4a and b), we only analyzed virions with up to 90 CHK-152 bound. These fell within a normally distributed portion of the population (Figure 4– Figure supplement 4b), in contrast with the lognormally distributed tail, and comprised 75% of the total number of virions observed.

Simulations. Numerical simulations were performed in Matlab. A grid of spikes was defined per Figure 6– Figure supplement 1b, where patch sizes from 12 to 40 (half a virion) were considered. Each spike contained 3 epitopes, and a specified number of inhibitors was bound randomly across all epitopes. This number of antibodies, or the related quantity epitope occupancy (number of antibodies divided by number of epitopes), was varied. Statistics were obtained for 10 000 virions. The number of unbound spikes within the contact patch was counted separately, and in the context of the defined 5- and 6-rings in Figure 6– Figure supplement 1b. The extent of fusion was defined as the fraction of virions that had at minimum one 5- or 6-ring with NH unbound spikes as shown in Figure 6c. To facilitate comparison with the numerical model (Figure 6e), the data was scaled to take into account dissociation. Effective number of CHK-152 bound: the average number of CHK-152 over non-fusing virions was averaged over time weighted by the number of unfused virions. This is therefore a measure for the average number of CHK-152 a fusing virion had bound during the time it took to fuse. Relative extent of fusion: the extent of fusion in the presence of CHK-152 was divided by the extent of fusion without antibody. The relative extent of fusion therefore is corrected for virions that were never able to fuse, and for the pH variability of the fusion extent.

## Acknowledgements

The authors wish to thank Dr Andres Merits and Dr Michael Diamond for their kind gifts of materials. We are grateful for technical assistance by Dr Viktor Krasnikov and the University of Groningen Workshop, and for the use of facilities of Dr E. Verpoorte with assistance of Patty Mulder.

## Supplements

**Figure 4– Table supplement 1.**
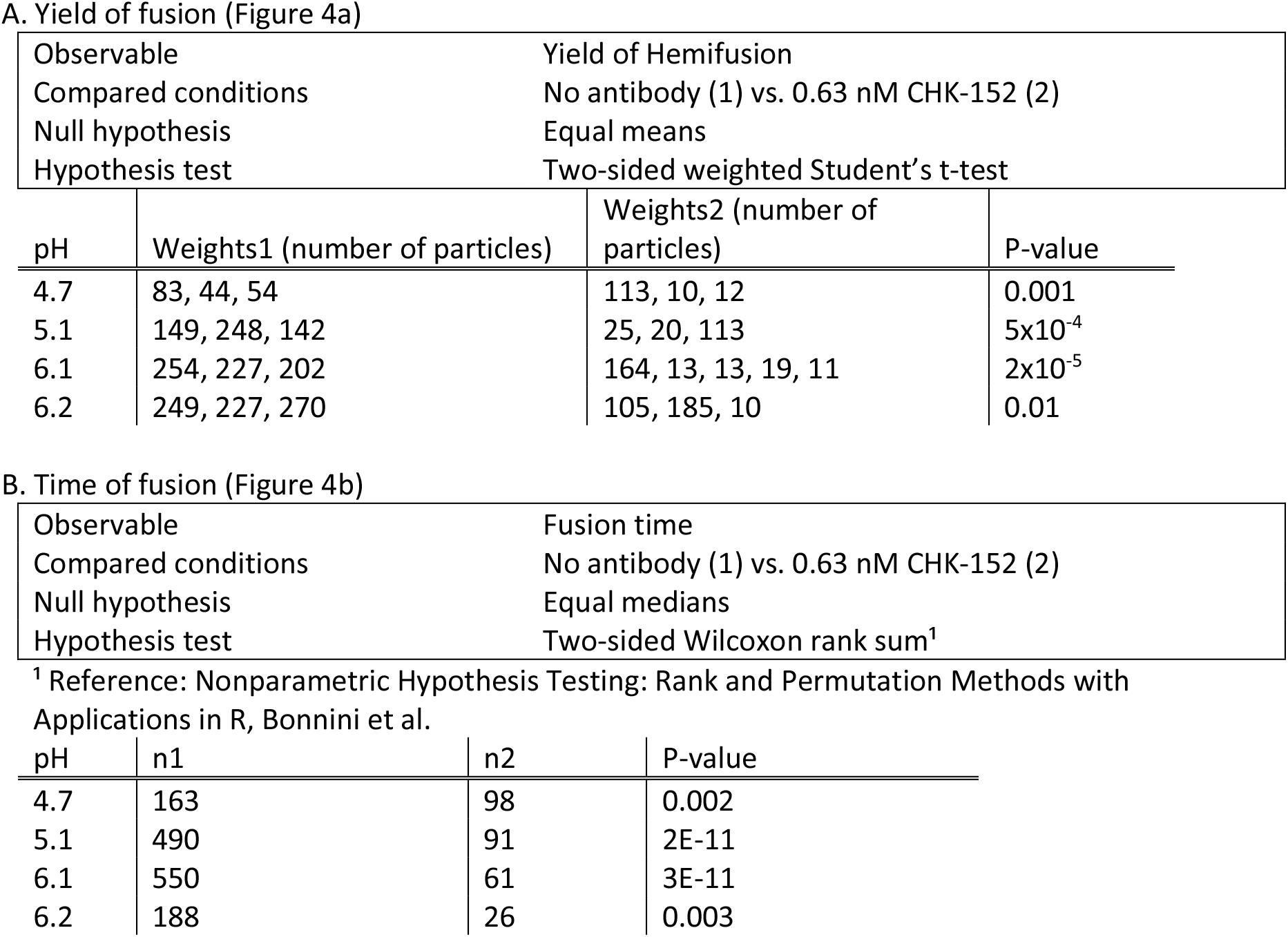
Significance testing.

| Observable |  | Yield of Hemifusion |  |
| --- | --- | --- | --- |
| Compared conditions |  | No antibody (1) vs. 0.63 nM CHK-152 (2) |  |
| Null hypothesis |  | Equal means |  |
| Hypothesis test |  | Two-sided weighted Student's t-test |  |
| pH | Weights1 (number of particles) | Weights2 (number of particles) | P-value |
| 4.7 | 83, 44, 54 | 113, 10, 12 | 0.001 |
| 5.1 | 149, 248, 142 | 25, 20, 113 | 5x10<sup>-4</sup> |
| 6.1 | 254, 227, 202 | 164, 13, 13, 19, 11 | 2x10<sup>-5</sup> |
| 6.2 | 249, 227, 270 | 105, 185, 10 | 0.01 |

B. Time of fusion (Figure 4b)
|  |  |  |
| --- | --- | --- |
| Observable |  | Fusion time |
| Compared conditions |  | No antibody (1) vs. 0.63 nM CHK-152 (2) |
| Null hypothesis |  | Equal medians |
| Hypothesis test |  | Two-sided Wilcoxon rank sum<sup>1</sup> |

<sup>1</sup> Reference: Nonparametric Hypothesis Testing: Rank and Permutation Methods with Applications in R, Bonnini et al.

| pH | n1 | n2 | P-value |
| --- | --- | --- | --- |
| 4.7 | 163 | 98 | 0.002 |
| 5.1 | 490 | 91 | 2E-11 |
| 6.1 | 550 | 61 | 3E-11 |
| 6.2 | 188 | 26 | 0.003 |

**Figure 4– Figure supplement 1.**
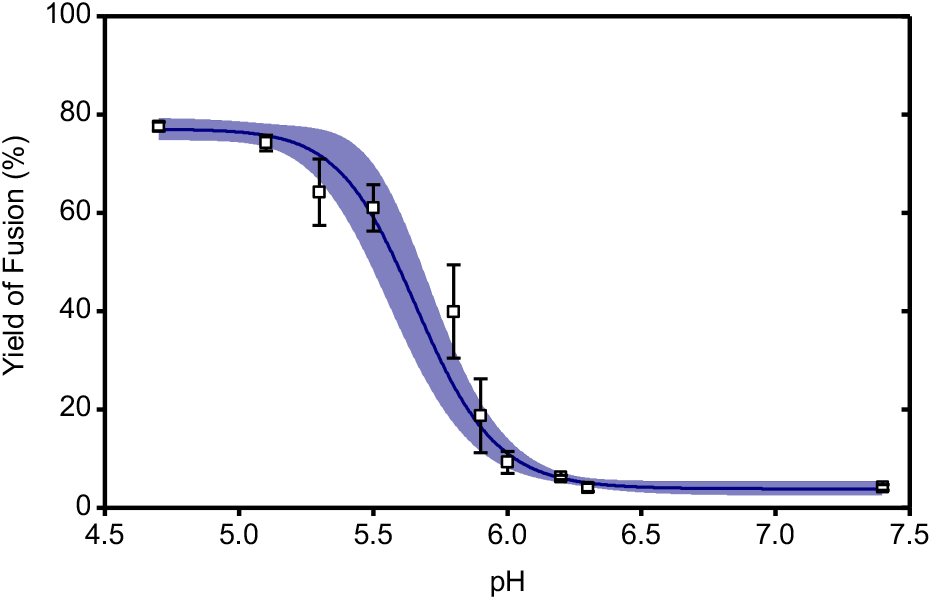
pH-dependence of the extent of fusion in a liposomal fusion assay for the CHIKV LR2006 OPY1 strain. Pyrene-labeled viruses were mixed with liposomes and acidified to the indicated pH at 37 °C. The yield of fusion was determined as the amount of fluorescence detected at 60 s relative to full dilution of the pyrene probe by detergent (see Methods). A logistic curve was fitted to the data, 95% confidence intervals indicated.

**Figure 4– Figure supplement 2.**
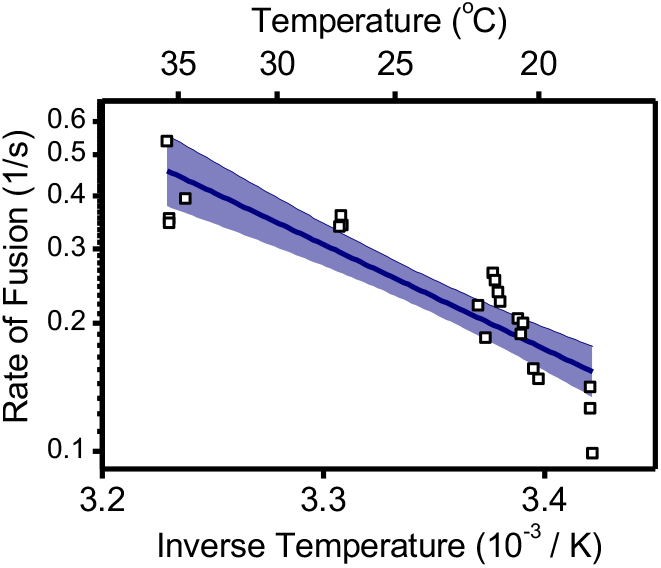
Arrhenius plot of the rate of fusion in a bulk liposomal fusion assay versus the inverse temperature. Pyrene-labeled viruses were mixed with liposomes and acidified to pH 5.1 at the specified temperature. The rate of fusion was determined as the inverse of the time to reach 50% of the maximum fusion extent (see Methods). *N*=21 trials. Linear fit with 95% confidence interval indicated.

**Figure 4– Figure supplement 3.**
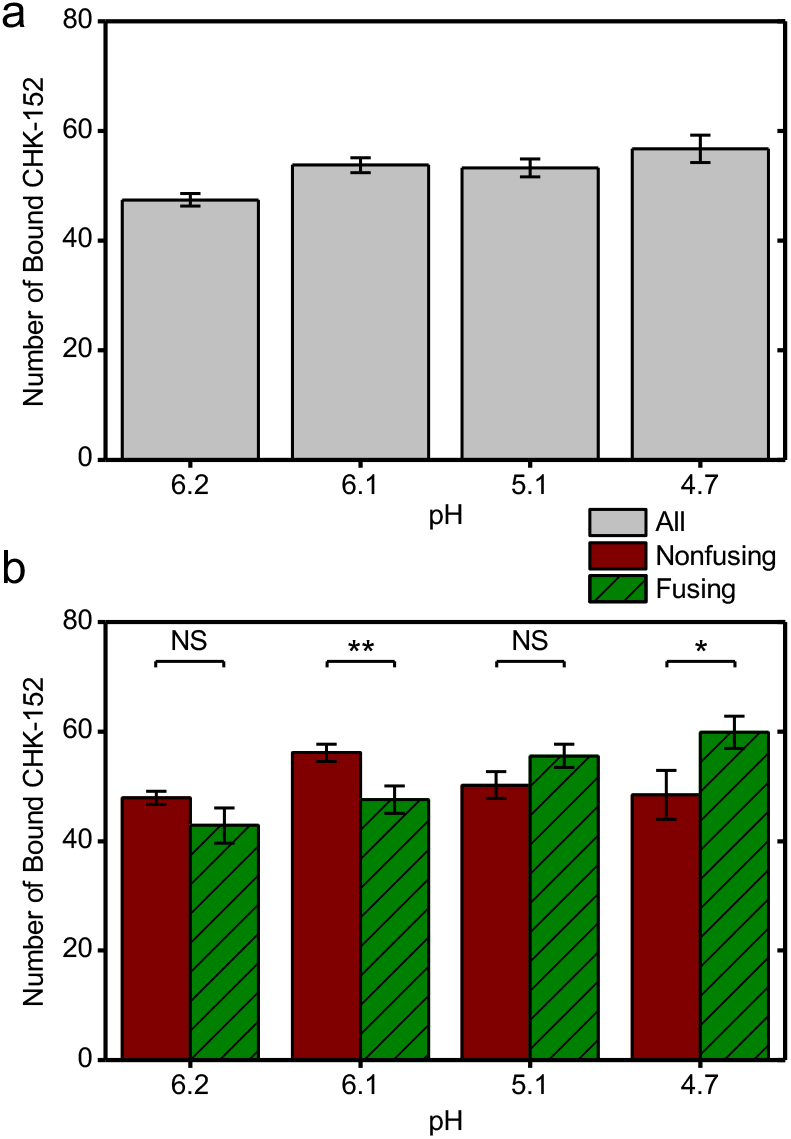
Number of CHK-152 bound at the start of the experiment per pH condition. As described in the Methods, the number of CHK-152 per virion was determined in the single-particle assay at *t* = 0. These numbers are here shown as means for different subsets of all virions. (a) All virions: number of CHK-152 bound at start for all particles taken together. (b) The data as in panel a split into the subpopulations of viruses that fuse and those that do not. Significances determined by t-test. * p<0.05, ** p<0.01.

**Figure 4– Figure supplement 4.**
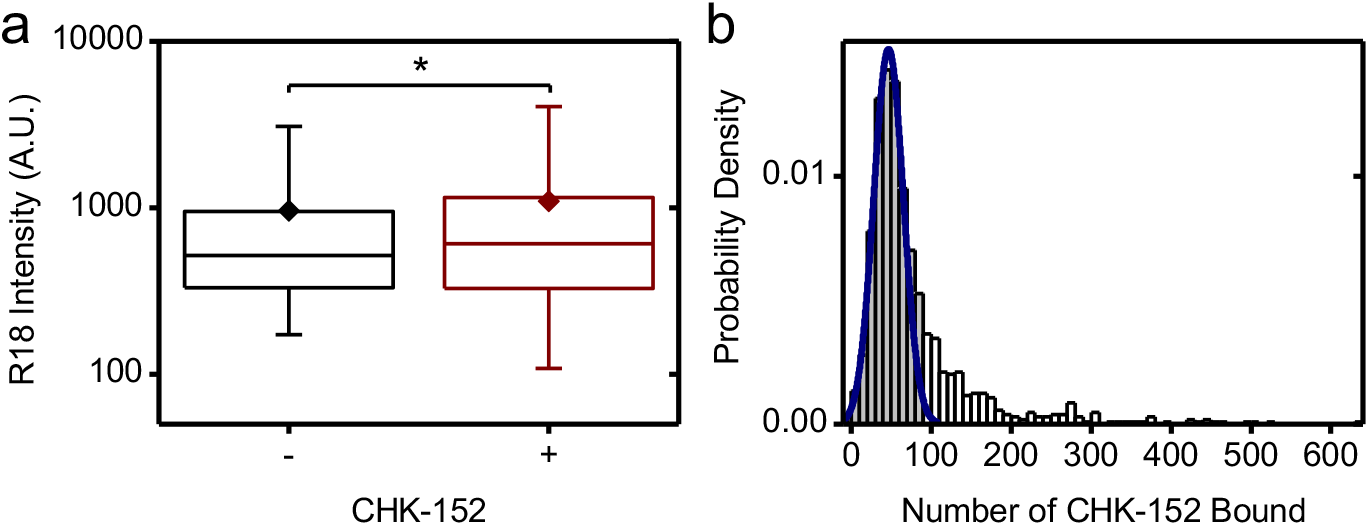
CHK-152-induced virion aggregation. a) For virions docked to the planar bilayer at pH 7.4 the R18 intensity was determined and is shown on a log scale. Conditions: -, without CHK-152 and +, with CHK-152 pre-incubation. Significance determined by t-test on the means, *n*-=2149 and *n*+=1042 virions. Means, diamonds; box plot shows 5%-Q1-median-Q3-95% intervals. b) Similarly, single-virion CHK-152 counts were determined at pH 7.4 and are shown in a histogram. Particles falling within the fitted Gaussian were selected for further analysis: the 75% of the data points with a CHK-152 count of up to 90 per virion.

**Figure 4– Movie Supplement 1.**
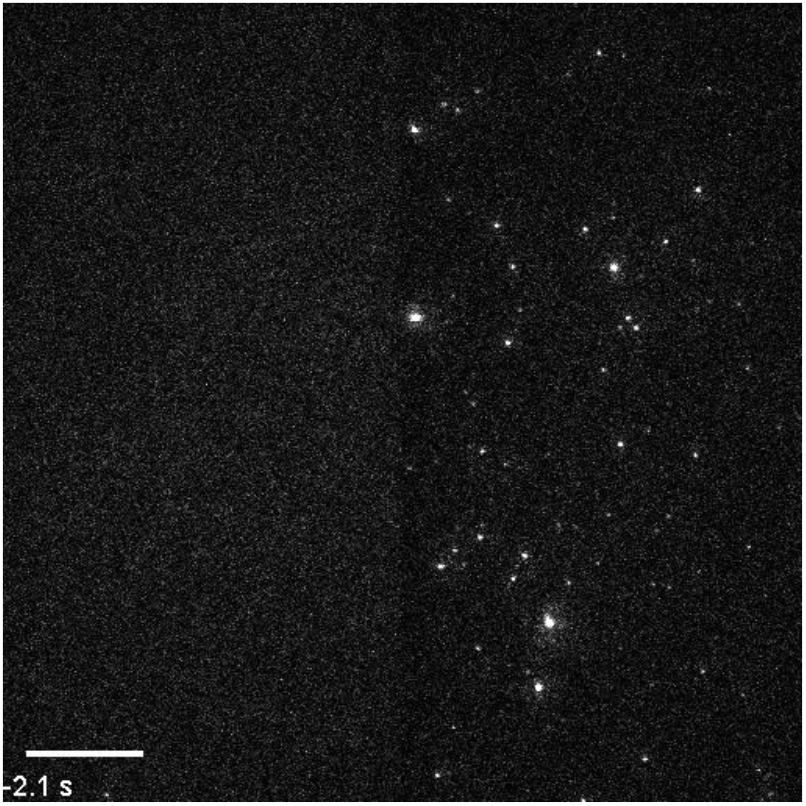
Single-particle CHIKV fusion at pH 4.7 without CHK-152. Scale bar 20 μm. Realtime timelapse. All timelapses

**Figure 4– Movie Supplement 2.**
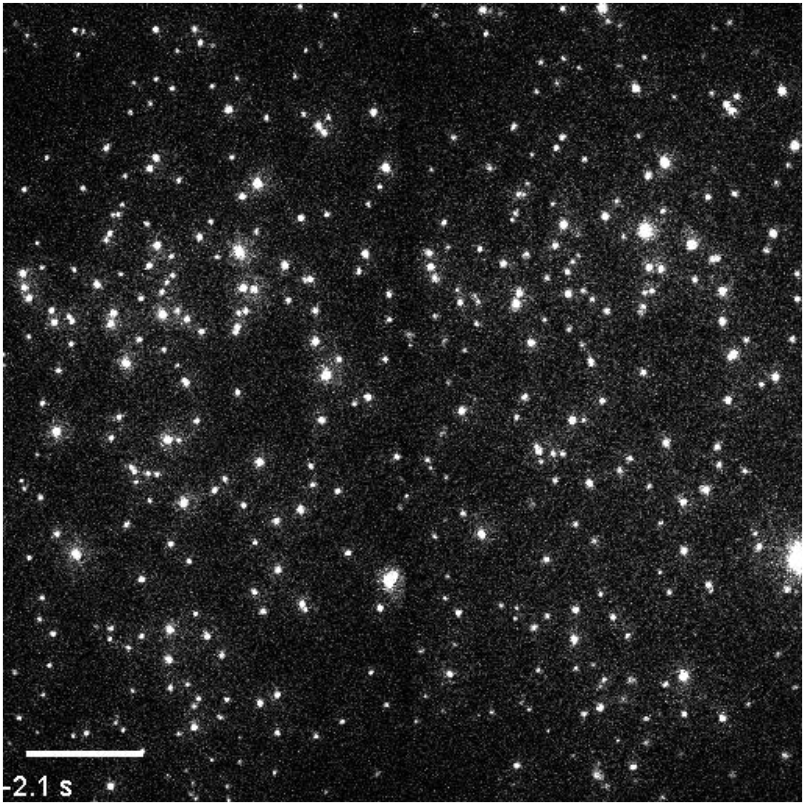
Timelapse of single-particle CHIKV fusion at pH 4.7 with CHK-152. Scale bar 20 μm. Realtime timelapse.

**Figure 4– Movie Supplement 3.**
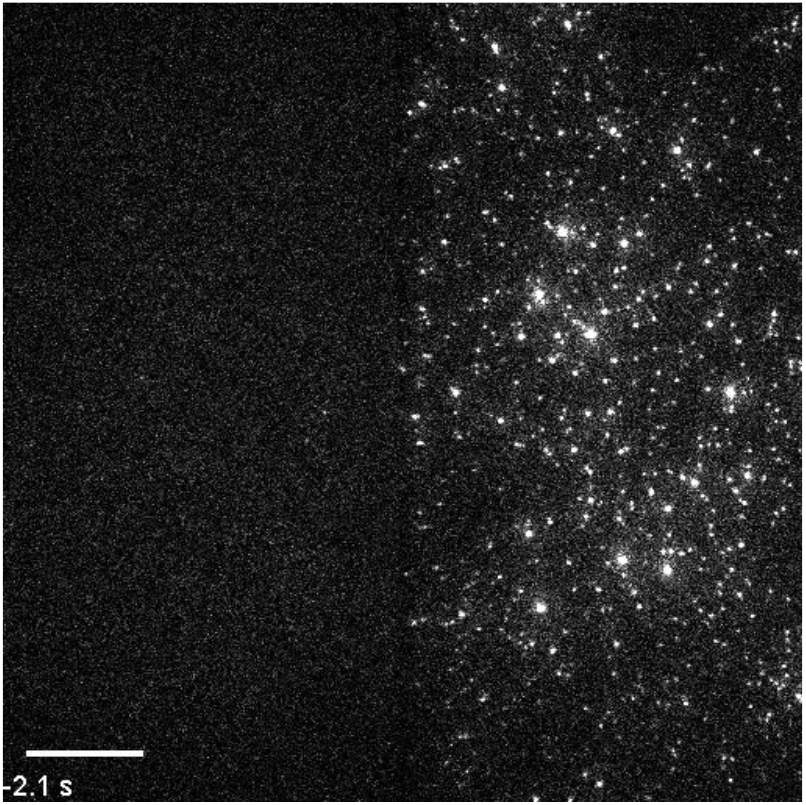
Timelapse of single-particle CHIKV fusion at pH 5.1 without CHK-152. Scale bar 20 μm. Realtime timelapse.

**Figure 4– Movie Supplement 4.**
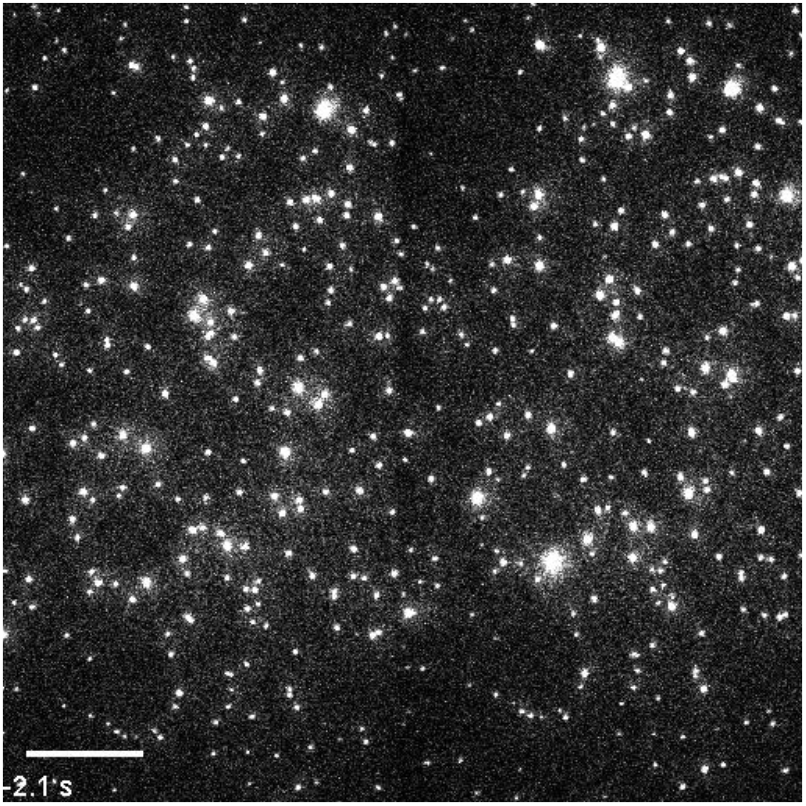
Timelapse of single-particle CHIKV fusion at pH 5.1 with CHK-152. Scale bar 20 μm. Realtime timelapse.

**Figure 4– Movie Supplement 5.**
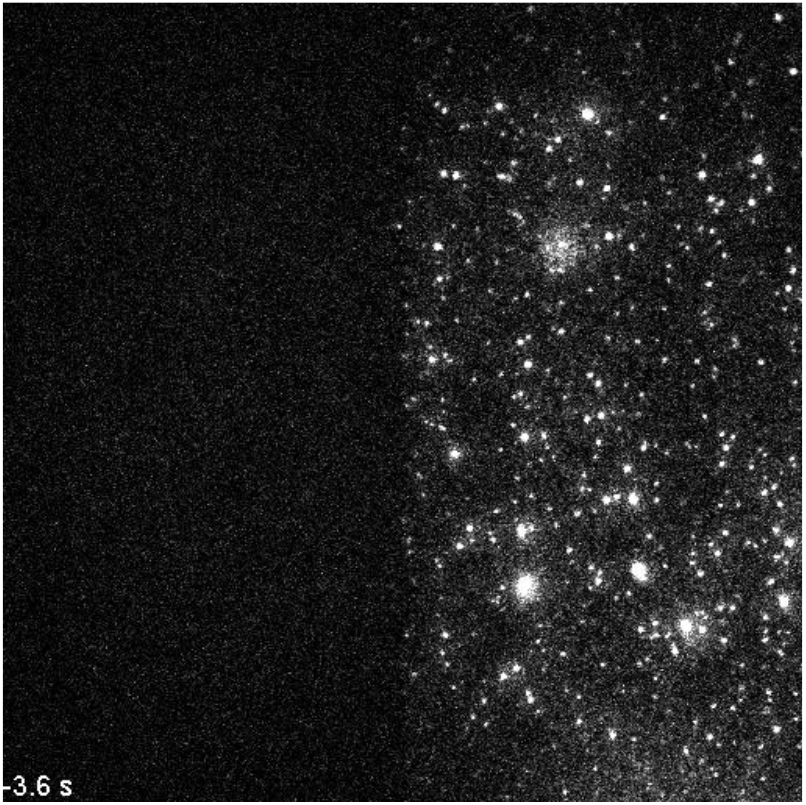
Timelapse of single-particle CHIKV fusion at pH 6.1 without CHK-152. Scale bar 20 μm. Timelapse at 3x speed.

**Figure 4– Movie Supplement 6.**
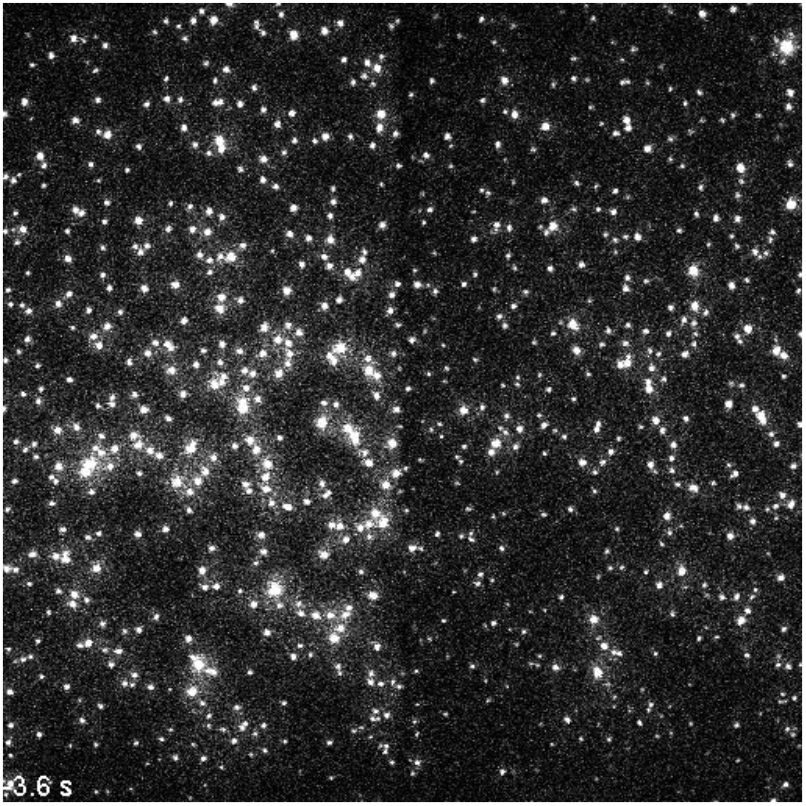
Timelapse of single-particle CHIKV fusion at pH 6.1 with CHK-152. Scale bar 20 μm. Timelapse at 3x speed.

**Figure 4– Movie Supplement 7.**
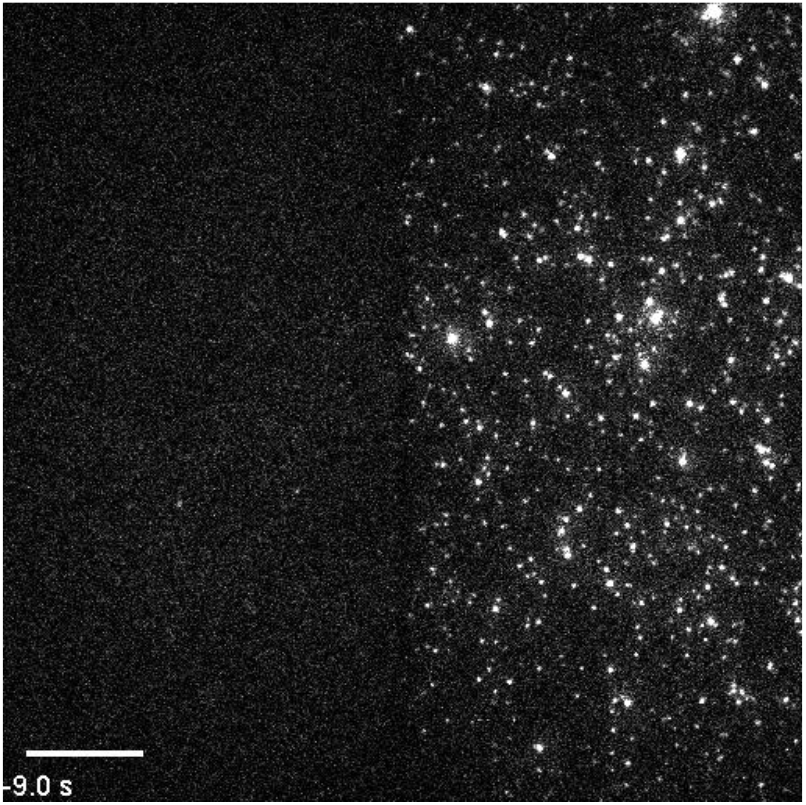
Timelapse of single-particle CHIKV fusion at pH 6.2 without CHK-152. Scale bar 20 μm. Timelapse at 5x speed.

**Figure 4– Movie Supplement 8.**
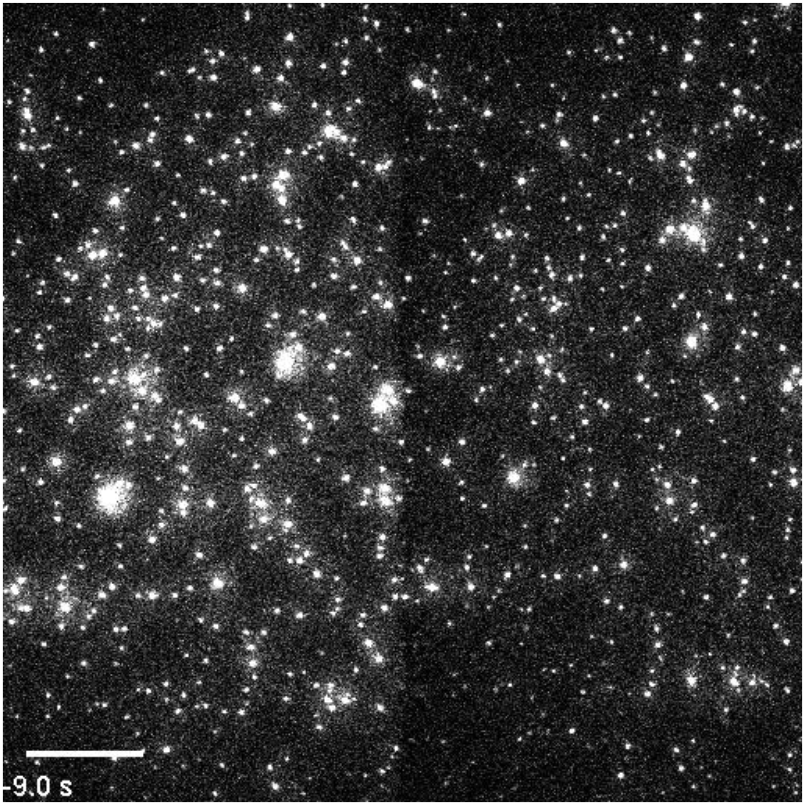
Timelapse of single-particle CHIKV fusion at pH 6.2 with CHK-152. Scale bar 20 μm. Timelapse at 5x speed.

**Figure 5– Table supplement 1.** Fitting functions used and resulting fit parameters.

| Parameter: | pH: | 6.2 | 6.1 |
| --- | --- | --- | --- |
| Baseline $y0$ | | $45.48 \pm 0.03$ | $41.45 \pm 0.08$ |
| Amplitude $A1$ | | $1.5 \pm 0.1$ | $2.05 \pm 0.08$ |
| Time scale $t1$ | | $8 \pm 1$ | $36 \pm 4$ |

Fit function: $y = A1 \cdot \exp(-x/t1) + A2 \cdot \exp(-x/t2) + y0$
| Parameter: | pH: | 5.1 | 4.7 |
| --- | --- | --- | --- |
| Baseline $y0$ | | $6.7 \pm 0.2$ | $3.83 \pm 0.03$ |
| Amplitude $A1$ | | $27 \pm 2$ | $4.9 \pm 0.3$ |
| Time scale $t1$ | | $59 \pm 3$ | $26 \pm 2$ |
| Amplitude $A2$ | | $12 \pm 2$ | $38 \pm 2$ |
| Time scale $t2$ | | $17 \pm 2$ | $3.4 \pm 0.2$ |

**Figure 5– Figure supplement 1.**
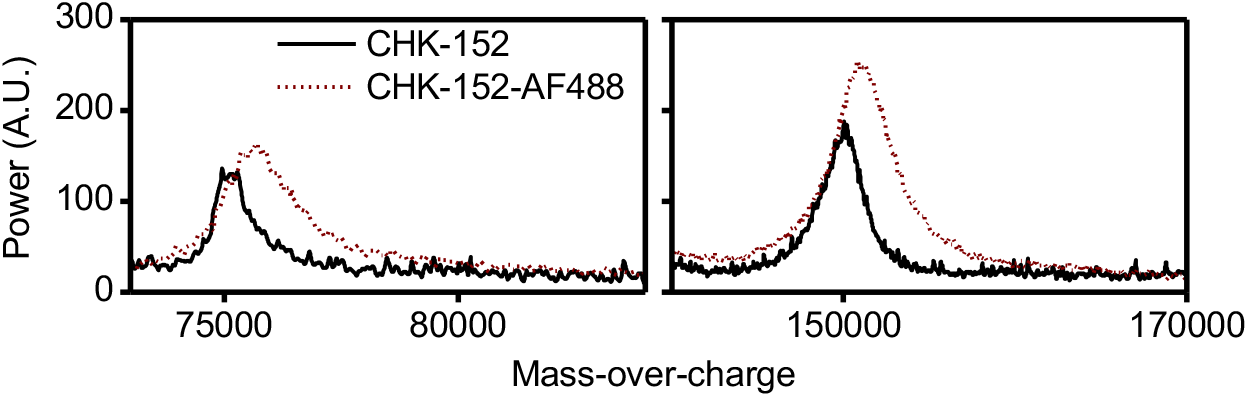
MALDI spectra of labeled and unlabeled CHK-152. Spectra were obtained with antibody dialyzed to 150 mM ammonium acetate. The AlexaFluor488 dye had a molecular weight of about 700.

**Figure 5– Figure supplement 2.**
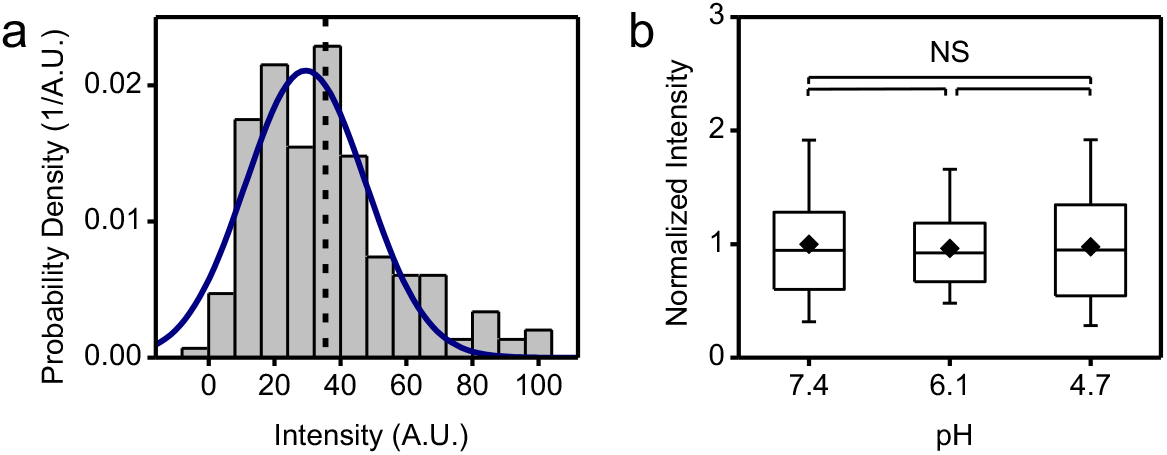
Labeled CHK-152 intensity determination. (a) Single AF488-labeled CHK-152 were flown into the flow cell and absorbed aspecifically to the cover glass at pH 7.4. Imaging conditions as for a fusion experiment were then used to extract the single CHK-152-AF488 intensity. The histogram of intensities is shown for n=186 spots. Solid line is a Gaussian fit, dashed line shows mean value. (b) With conditions as in panel a, the intensities of CHK-152-AF488 versus pH are shown, normalized to mean pH 7.4 intensity. Significances from t-test, *n* = 65,47,58 spots respectively. Means, diamonds; box plot shows 5%-Q1-median-Q3-95% intervals.

**Figure 5– Figure supplement 3.**
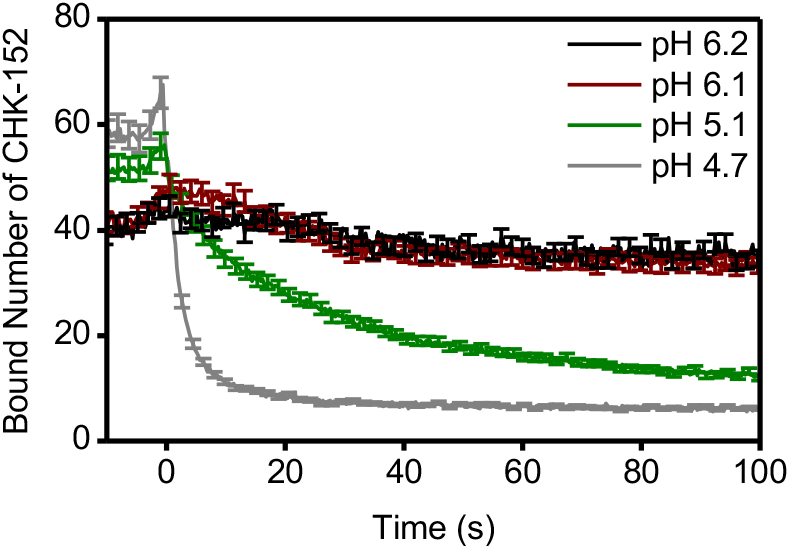
Bound number of CHK-152 averaged for all fusing particles over time. In the single-particle assay, the fluorescence intensity of virions was tracked over time and converted to absolute number of CHK-152 bound (Methods). The average number of CHK-152 bound for fusing virions is shown over time. Increase of signal towards *t* = 0 due to rolling and arrest of virus particles. One out of every five error bars (sem) shown to reduce visual clutter.

**Figure 6– Figure supplement 1.**
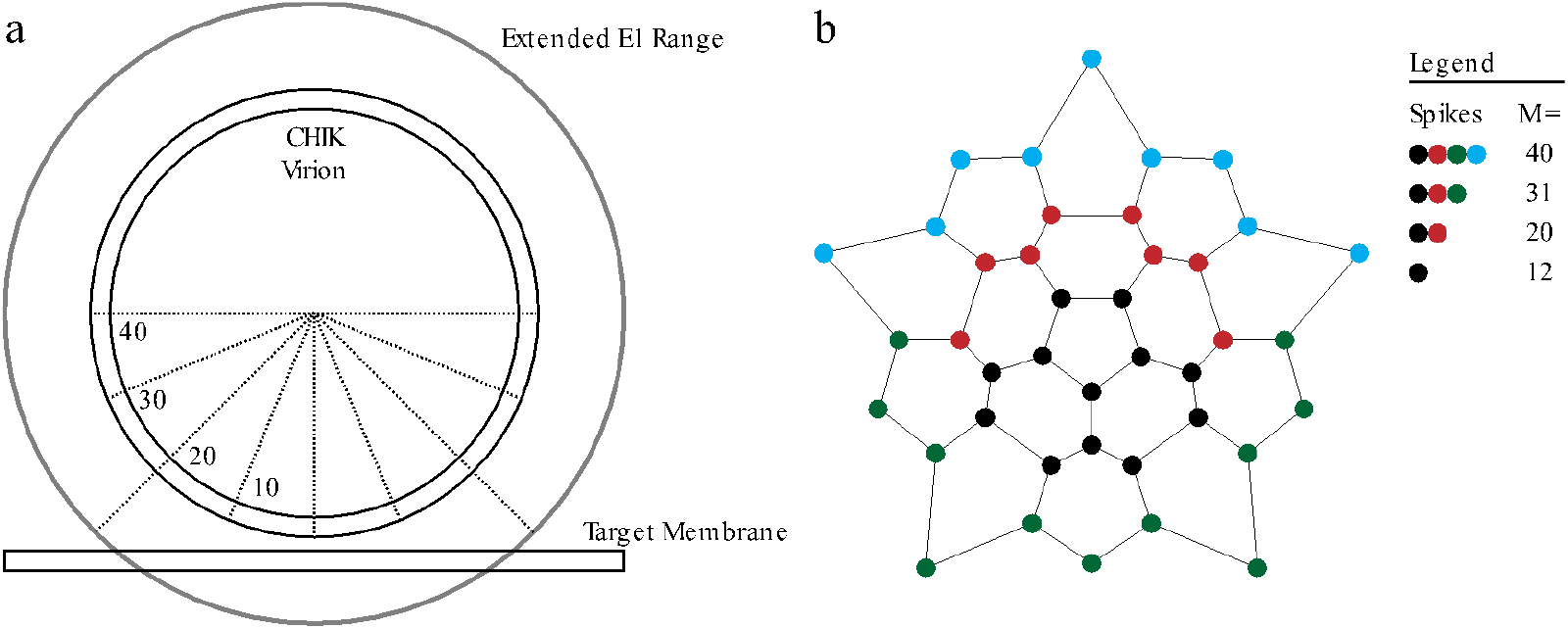
Patch size considerations. (a) Schematic diagram of the number of spikes that fall within range of the contact patch (delineated by dotted lines) facing the target membrane. Virion of 65 nm diameter and E1 proteins of 13 nm length assumed (approximate range shown in grey). The number of spikes is shown, that fits on the relative fraction of the viral surface indicated. In total the virion comprises 80 spikes. (b) Layout of the surface grid of spikes of one half of a CHIK virion. The lines indicate the connections that make rings. The different contact patch sizes are indicated by color, cumulatively: *M* = 12 (black), *M* = 20 (black+red), *M* = 31 (black+red+green), *M* = 40 (black+red+green+blue).

**Figure 6– Figure supplement 2.**
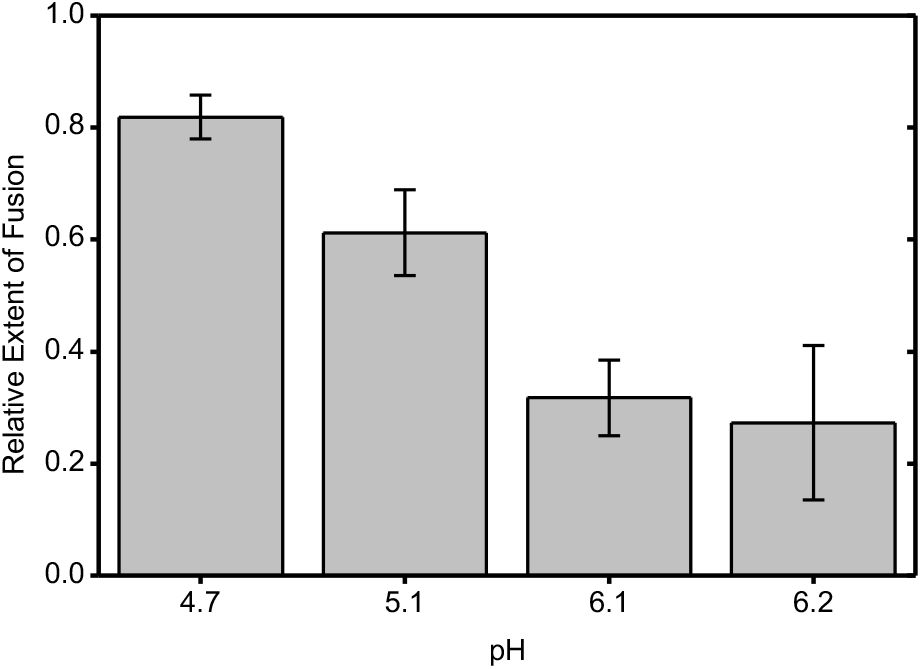
Relative extent of fusion with CHK-152 for each pH point. The relative extent of fusion was calculated as the ratio of the extents of fusion of the antibody and no-antibody conditions (both from **Figure 4**). Sem was propagated accordingly.

**Figure 6– Figure supplement 3.**
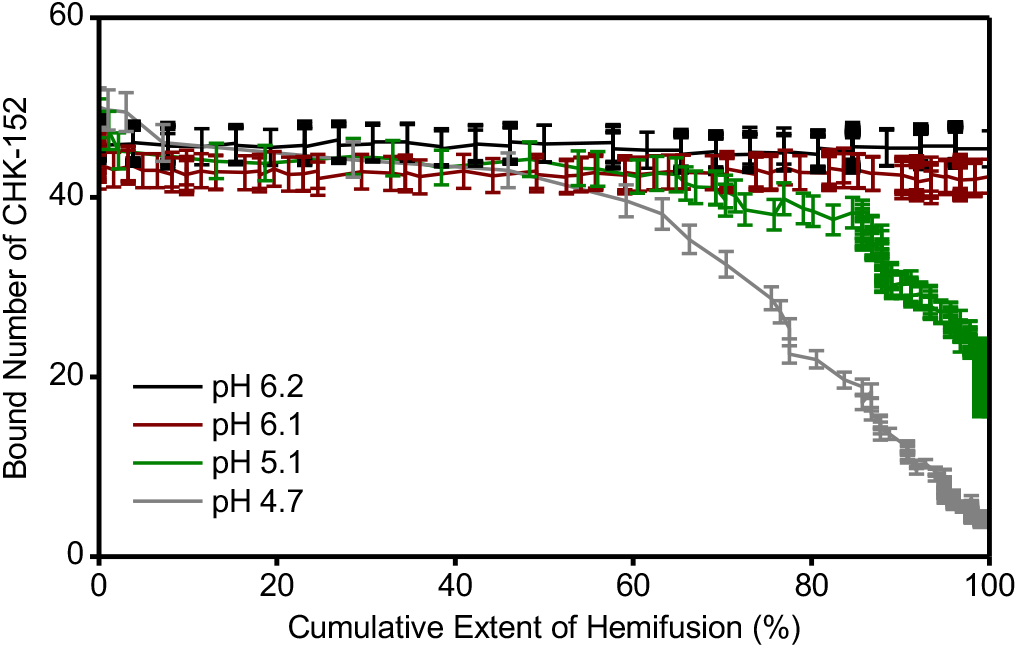
Correlation of the mean number of bound CHK-152 versus the cumulative extent of fusion. Both the extent of fusion and CHK-152 number were determined over time for individual virions and then averaged. The two readouts are here plotted against each other for each time point. As is visible in Figure 5, at pH 6.2 and 6.1 only a small number of CHK-152 dissociate, whereas at pH 5.1 and 4.7 dissociation occurs. The graph shows that at pH 5.1 and 4.7 only late-fusing virions, with respect to the whole fusing population, have lost large numbers of CHK-152 at the moment of fusion.

**Figure 6– Figure supplement 4.**
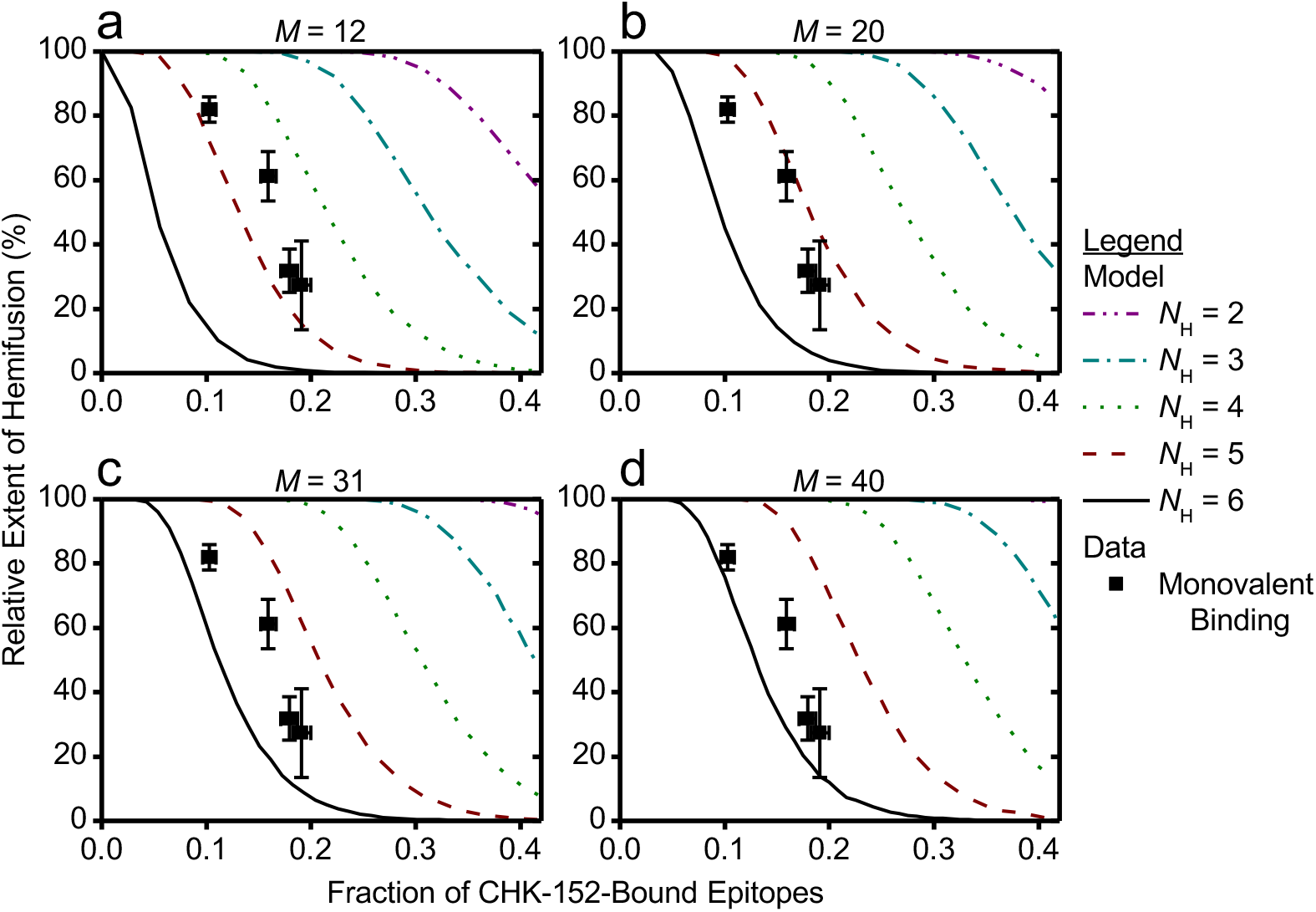
Simulation and data compared for different patch sizes, assuming monovalent CHK-152 binding. Like in Figure 6, for 10 000 virions CHK-152 was randomly bound and the relative extent of fusion was determined as the fraction of virions having available *N*H free spikes in a ring. The extents of fusion from the simulations are shown as lines versus the fraction of CHK-152-bound epitopes on the viral surface. Line legends are as shown in Figure 6c: *N*H = 3,4,5,6 are indicated by dash-dotted, dotted, dashed and a solid line respectively. The data points (squares) shown are the same for every graph and are equal to that of Figure 6e. The simulation was adapted to assume a contact patch of *M* = 12,20,31,40 spikes as indicated above the graphs.

**Figure 6– Figure supplement 5.**
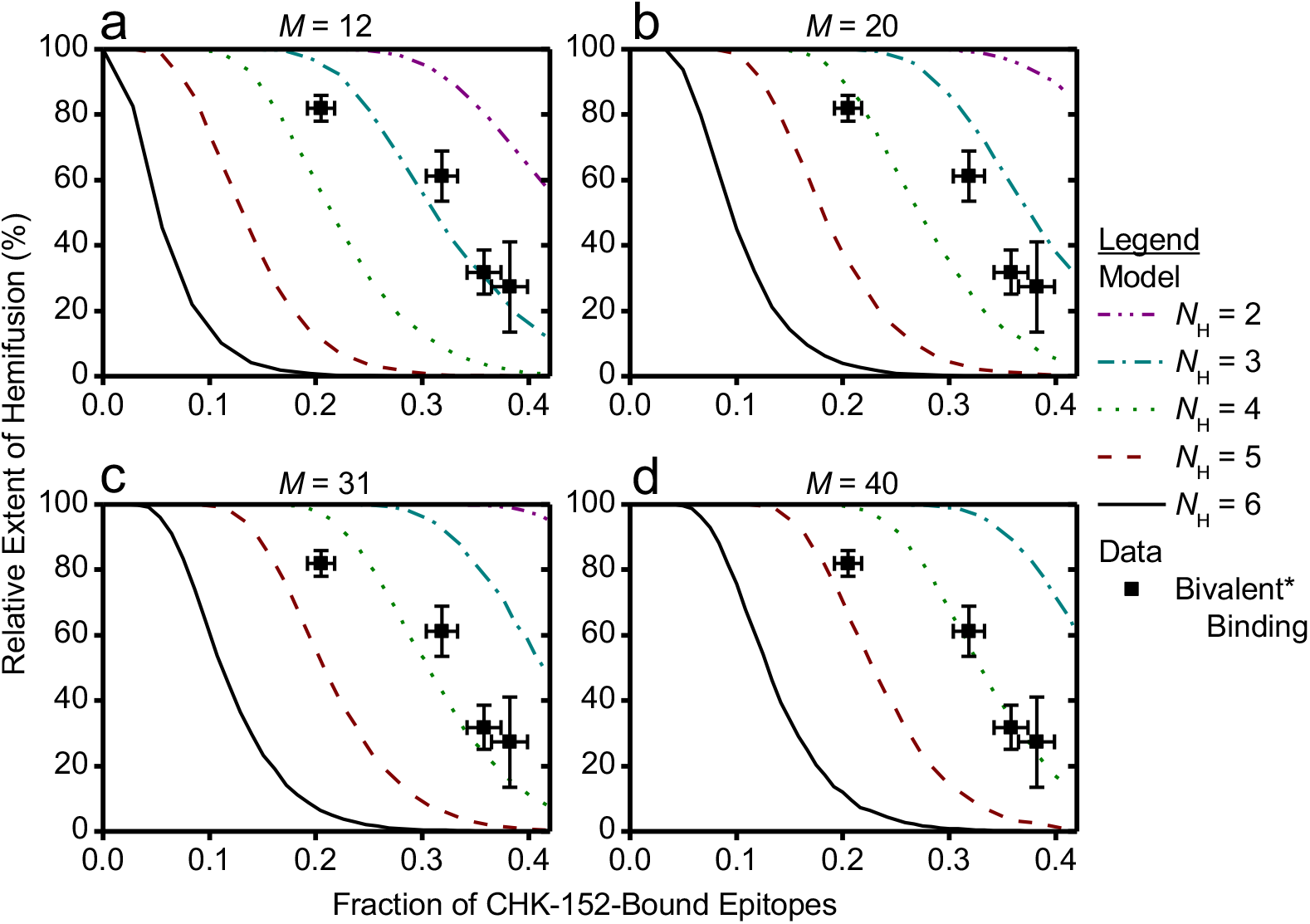
Simulation and data compared for different patch sizes, assuming bivalent CHK-152 binding. Simulation, legend and data like Figure 6– Figure supplement 4, but assuming bivalent* binding of CHK-152. This was modeled by binding double the amount of Fabs.

## References

Ahn, A., Gibbons, D.L. & Kielian, M. 2002, “The fusion peptide of Semliki Forest virus associates with sterol-rich membrane domains”, Journal of virology, vol. 76, no. 7, pp. 3267–3275.

Ashbrook, A.W., Burrack, K.S., Silva, L.A., Montgomery, S.A., Heise, M.T., Morrison, T.E. & Dermody, T.S. 2014, “Residue 82 of the Chikungunya virus E2 attachment protein modulates viral dissemination and arthritis in mice”, Journal of virology, vol. 88, no. 21, pp. 12180–12192 doi:10.1128/JVI.01672-14 [doi].

Bernard, E., Solignat, M., Gay, B., Chazal, N., Higgs, S., Devaux, C. & Briant, L. 2010, “Endocytosis of chikungunya virus into mammalian cells: role of clathrin and early endosomal compartments”, PloS one, vol. 5, no. 7, pp. e11479 doi:10.1371/journal.pone.0011479.

Bonizzoni, M., Gasperi, G., Chen, X. & James, A.A. 2013, “The invasive mosquito species Aedes albopictus: current knowledge and future perspectives”, Trends in parasitology, vol. 29, no. 9, pp. 460–468 doi:10.1016/j.pt.2013.07.003 [doi].

Bron, R., Wahlberg, J.M., Garoff, H. & Wilschut, J. 1993, “Membrane fusion of Semliki Forest virus in a model system: correlation between fusion kinetics and structural changes in the envelope glycoprotein”, The EMBO journal, vol. 12, no. 2, pp. 693–701.

Cao, S. & Zhang, W. 2013, “Characterization of an early-stage fusion intermediate of Sindbis virus using cryoelectron microscopy”, Proceedings of the National Academy of Sciences of the United States of America, vol. 110, no. 33, pp. 13362–13367 doi:10.1073/pnas.1301911110 [doi].

Centers for Disease Control and Prevention (CDC), Geographic Distribution of Chikungunya Virus. Available online: https://www.cdc.gov/chikungunya/geo/index.html.

Chao, L.H., Klein, D.E., Schmidt, A.G., Pena, J.M. & Harrison, S.C. 2014, “Sequential conformational rearrangements in flavivirus membrane fusion”, Elife, vol. 3, pp. e04389 doi:10.7554/eLife.04389.

Chernomordik, L.V. & Kozlov, M.M. 2008, “Mechanics of membrane fusion”, Nature Structural & Molecular Biology, vol. 15, no. 7, pp. 675–683 doi:10.1038/nsmb.1455.

Clayton, A.M. 2016, “Monoclonal Antibodies as Prophylactic and Therapeutic Agents Against Chikungunya Virus”, The Journal of infectious diseases, vol. 214, no. suppl 5, pp. S506–S509 doi:10.1093/infdis/jiw324.

Floyd, D.L., Ragains, J.R., Skehel, J.J., Harrison, S.C. & van Oijen, A.M. 2008, “Single-particle kinetics of influenza virus membrane fusion”, Proceedings of the National Academy of Sciences of the United States of America, vol. 105, no. 40, pp. 15382–15387 doi:10.1073/pnas.0807771105.

Fox, J.M., Long, F., Edeling, M.A., Lin, H., van Duijl-Richter, M.K.S., Fong, R.H., Kahle, K.M., Smit, J.M., Jin, J., Simmons, G., Doranz, B.J., Crowe, J.E., Jr, Fremont, D.H., Rossmann, M.G. & Diamond, M.S. 2015, “Broadly Neutralizing Alphavirus Antibodies Bind an Epitope on E2 and Inhibit Entry and Egress”, Cell, vol. 163, no. 5, pp. 1095–1107 doi:10.1016/j.cell.2015.10.050.

Gibbons, D.L., Erk, I., Reilly, B., Navaza, J., Kielian, M., Rey, F.A. & Lepault, J. 2003, “Visualization of the target-membrane-inserted fusion protein of Semliki Forest virus by combined electron microscopy and crystallography”, Cell, vol. 114, no. 5, pp. 573–583 doi:10.1016/S0092-8674(03)00683-4.

Gibbons, D.L., Vaney, M.C., Roussel, A., Vigouroux, A., Reilly, B., Lepault, J., Kielian, M. & Rey, F.A. 2004, “Conformational change and protein-protein interactions of the fusion protein of Semliki Forest virus”, Nature, vol. 427, no. 6972, pp. 320–325 doi:10.1038/nature02239 [doi].

Harrison, S.C. 2015, “Viral membrane fusion”, Virology, vol. 479, pp. 498–507 doi:10.1016/j.virol.2015.03.043.

Hoornweg, T.E., van Duijl-Richter, M.K., Ayala Nunez, N.V., Albulescu, I.C., van Hemert, M.J. & Smit, J.M. 2016, “Dynamics of Chikungunya Virus Cell Entry Unraveled by Single-Virus Tracking in Living Cells”, Journal of virology, vol. 90, no. 9, pp. 4745–4756 doi:10.1128/JVI.03184-15 [doi].

Ivanovic, T., Choi, J.L., Whelan, S.P.J., van Oijen, A.M. & Harrison, S.C. 2013, “Influenza virus membrane fusion by cooperative fold-back of stochastically induced hemagglutinin intermediates”, eLife, vol. 2, no. 0, pp. e00333 doi:10.7554/eLife.00333.

Ivanovic, T. & Harrison, S.C. 2015, “Distinct functional determinants of influenza hemagglutinin-mediated membrane fusion”, eLife, vol. 4, pp. e11009 doi:10.7554/eLife.11009.

Jin, J., Liss, N.M., Chen, D.H., Liao, M., Fox, J.M., Shimak, R.M., Fong, R.H., Chafets, D., Bakkour, S., Keating, S., Fomin, M.E., Muench, M.O., Sherman, M.B., Doranz, B.J., Diamond, M.S. & Simmons, G. 2015, “Neutralizing Monoclonal Antibodies Block Chikungunya Virus Entry and Release by Targeting an Epitope Critical to Viral Pathogenesis”, Cell reports, vol. 13, no. 11, pp. 2553–2564 doi:10.1016/j.celrep.2015.11.043.

Jose, J., Snyder, J.E. & Kuhn, R.J. 2009, “A structural and functional perspective of alphavirus replication and assembly”, Future microbiology,vol. 4, no. 7, pp. 837–856 doi:10.2217/fmb.09.59 [doi].

Kaufmann, B., Vogt, M.R., Goudsmit, J., Holdaway, H.A., Aksyuk, A.A., Chipman, P.R., Kuhn, R.J., Diamond, M.S. & Rossmann, M.G. 2010, “Neutralization of West Nile virus by cross-linking of its surface proteins with Fab fragments of the human monoclonal antibody CR4354”, Proceedings of the National Academy of Sciences of the United States of America, vol. 107, no. 44, pp. 18950–18955 doi:10.1073/pnas.1011036107 [doi].

Kielian, M. & Helenius, A. 1985, “pH-induced alterations in the fusogenic spike protein of Semliki Forest virus.“, The Journal of cell biology, vol. 101, no. 6, pp. 2284–2291.

Kielian, M. 2014, “Mechanisms of virus membrane fusion proteins”, Annual Review of Virology, vol. 1, pp. 171–189 doi:10.1146/annurev-virology-031413-085521.

Kim, I.S., Jenni, S., Stanifer, M.L., Roth, E., Whelan, S.P.J., van Oijen, A.M. & Harrison, S.C. 2017, “Mechanism of membrane fusion induced by vesicular stomatitis virus G protein”, Proceedings of the National Academy of Sciences of the United States of America, vol. 114, no. 1, pp. E28–E36 doi:10.1073/pnas.1618883114.

Klimjack, M.R., Jeffrey, S. & Kielian, M. 1994, “Membrane and protein interactions of a soluble form of the Semliki Forest virus fusion protein”, Journal of virology, vol. 68, no. 11, pp. 6940–6946.

Kolter, T. & Sandhoff, K. 2010, “Lysosomal degradation of membrane lipids”, FEBS letters, vol. 584, no. 9, pp. 1700–1712 doi:10.1016/j.febslet.2009.10.021 [doi].

Li, L., Jose, J., Xiang, Y., Kuhn, R.J. & Rossmann, M.G. 2010, “Structural changes of envelope proteins during alphavirus fusion”, Nature, vol. 468, no. 7324, pp. 705–708 doi:10.1038/nature09546 [doi].

Nieva, J.L., Bron, R., Corver, J. & Wilschut, J. 1994, “Membrane fusion of Semliki Forest virus requires sphingolipids in the target membrane”, The EMBO journal, vol. 13, no. 12, pp. 2797–2804.

Otterstrom, J.J., Brandenburg, B., Koldijk, M.H., Juraszek, J., Tang, C., Mashaghi, S., Kwaks, T., Goudsmit, J., Vogels, R., Friesen, R.H.E. & van Oijen, A.M. 2014, “Relating influenza virus membrane fusion kinetics to stoichiometry of neutralizing antibodies at the single-particle level”, Proceedings of the National Academy of Sciences of the United States of America, vol. 111, no. 48, pp. E5143–E5148 doi:10.1073/pnas.1411755111.

Pal, P., Dowd, K.A., Brien, J.D., Edeling, M.A., Gorlatov, S., Johnson, S., Lee, I., Akahata, W., Nabel, G.J., Richter, M.K., Smit, J.M., Fremont, D.H., Pierson, T.C., Heise, M.T. & Diamond, M.S. 2013, “Development of a highly protective combination monoclonal antibody therapy against Chikungunya virus”, PLoS pathogens, vol. 9, no. 4, pp. e1003312 doi:10.1371/journal.ppat.1003312 [doi].

Pal, P., Fox, J.M., Hawman, D.W., Huang, Y.J., Messaoudi, I., Kreklywich, C., Denton, M., Legasse, A.W., Smith, P.P., Johnson, S., Axthelm, M.K., Vanlandingham, D.L., Streblow, D.N., Higgs, S., Morrison, T.E. & Diamond, M.S. 2014, “Chikungunya viruses that escape monoclonal antibody therapy are clinically attenuated, stable, and not purified in mosquitoes”, Journal of virology,vol. 88, no. 15, pp. 8213–8226 doi:10.1128/JVI.01032-14 [doi].

Reiter, P., Fontenille, D. & Paupy, C. 2006, “Aedes albopictus as an epidemic vector of chikungunya virus: another emerging problem?”, The Lancet.Infectious diseases, vol. 6, no. 8, pp. 463–464 doi:10.1016/S1473-3099(06)70531-X.

Rueden, C.T., Schindelin, J., Hiner, M.C., DeZonia, B.E., Walter, A.E., Arena, E.T. & Eliceiri, K.W. 2017, “ImageJ2: ImageJ for the next generation of scientific image data”, BMC Bioinformatics, vol. 18, no. 1, pp. 529 doi:10.1186/s12859-017-1934-z.

Schindelin, J., Arganda-Carreras, I., Frise, E., Kaynig, V., Longair, M., Pietzsch, T., Preibisch, S., Rueden, C., Saalfeld, S., Schmid, B., Tinevez, J., White, D.J., Hartenstein, V., Eliceiri, K., Tomancak, P. & Cardona, A. 2012, “Fiji: an open-source platform for biological-image analysis”, Nature Methods, vol. 9, pp. 676.

Selvarajah, S., Sexton, N.R., Kahle, K.M., Fong, R.H., Mattia, K.A., Gardner, J., Lu, K., Liss, N.M., Salvador, B., Tucker, D.F., Barnes, T., Mabila, M., Zhou, X., Rossini, G., Rucker, J.B., Sanders, D.A., Suhrbier, A., Sambri, V., Michault, A., Muench, M.O., Doranz, B.J. & Simmons, G. 2013, “A neutralizing monoclonal antibody targeting the acid-sensitive region in chikungunya virus E2 protects from disease”, PLoS neglected tropical diseases, vol. 7, no. 9, pp. e2423 doi:10.1371/journal.pntd.0002423 [doi].

Smit, J.M., Bittman, R. & Wilschut, J. 1999, “Low-pH-dependent fusion of Sindbis virus with receptor-free cholesterol- and sphingolipid-containing liposomes”, Journal of virology, vol. 73, no. 10, pp. 8476–8484.

Smith, S.A., Silva, L.A., Fox, J.M., Flyak, A.I., Kose, N., Sapparapu, G., Khomadiak, S., Ashbrook, A.W., Kahle, K.M., Fong, R.H., Swayne, S., Doranz, B.J., McGee, C.E., Heise, M.T., Pal, P., Brien, J.D., Austin, S.K., Diamond, M.S., Dermody, T.S. & Crowe, J.E.,Jr 2015, “Isolation and Characterization of Broad and Ultrapotent Human Monoclonal Antibodies with Therapeutic Activity against Chikungunya Virus”, Cell host & microbe, vol. 18, no. 1, pp. 86–95 doi:10.1016/j.chom.2015.06.009 [doi].

Smith, T.J., Cheng, R.H., Olson, N.H., Peterson, P., Chase, E., Kuhn, R.J. & Baker, T.S. 1995, “Putative receptor binding sites on alphaviruses as visualized by cryoelectron microscopy”, Proceedings of the National Academy of Sciences of the United States of America, vol. 92, no. 23, pp. 10648–10652 doi:10.1073/pnas.92.23.10648.

Sun, S., Xiang, Y., Akahata, W., Holdaway, H., Pal, P., Zhang, X., Diamond, M.S., Nabel, G.J. & Rossmann, M.G. 2013, “Structural analyses at pseudo atomic resolution of Chikungunya virus and antibodies show mechanisms of neutralization”, eLife, vol. 2, pp. e00435 doi:10.7554/eLife.00435 [doi].

van Duijl-Richter, M.K.S. 2016, Dengue and Chikungunya virus: Cell entry mechanisms and the impact of antibodies on infectivity, University of Groningen http://hdl.handle.net/11370/7c4c710d-0bb2-4b84-8040-4795a14e9ee7.

van Duijl-Richter, M.K., Blijleven, J.S., van Oijen, A.M. & Smit, J.M. 2015, “Chikungunya virus fusion properties elucidated by single-particle and bulk approaches”, The Journal of general virology,vol. 96, no. 8, pp. 2122–2132 doi:10.1099/vir.0.000144 [doi].

van Meer, G., Voelker, D.R. & Feigenson, G.W. 2008, “Membrane lipids: where they are and how they behave”, Nature Reviews Molecular Cell Biology, vol. 9, no. 2, pp. 112–124 doi:10.1038/nrm2330.

Voss, J.E., Vaney, M.C., Duquerroy, S., Vonrhein, C., Girard-Blanc, C., Crublet, E., Thompson, A., Bricogne, G. & Rey, F.A. 2010, “Glycoprotein organization of Chikungunya virus particles revealed by X-ray crystallography”, Nature, vol. 468, no. 7324, pp. 709–712 doi:10.1038/nature09555 [doi].

Wahlberg, J.M., Boere, W.A. & Garoff, H. 1989, “The heterodimeric association between the membrane proteins of Semliki Forest virus changes its sensitivity to low pH during virus maturation”, Journal of virology, vol. 63, no. 12, pp. 4991–4997.

Wahlberg, J.M., Bron, R., Wilschut, J. & Garoff, H. 1992, “Membrane fusion of Semliki Forest virus involves homotrimers of the fusion protein”, Journal of virology, vol. 66, no. 12, pp. 7309–7318.

Weaver, S.C. & Forrester, N.L. 2015, “Chikungunya: Evolutionary history and recent epidemic spread”, Antiviral Research, vol. 120, pp. 32–39 doi:10.1016/j.antiviral.2015.04.016.

Zeng, X., Mukhopadhyay, S. & Brooks, C.L.,3rd 2015, “Residue-level resolution of alphavirus envelope protein interactions in pH-dependent fusion”, Proceedings of the National Academy of Sciences of the United States of America, vol. 112, no. 7, pp. 2034–2039 doi:10.1073/pnas.1414190112 [doi].

Zhang, Y. & Dudko, O.K. 2015, “Statistical mechanics of viral entry”, Physical Review Letters, vol. 114, no. 1, pp. 018104 doi:10.1103/PhysRevLett.114.018104.

Zheng, Y., Sanchez-San Martin, C., Qin, Z.L. & Kielian, M. 2011, “The domain I-domain III linker plays an important role in the fusogenic conformational change of the alphavirus membrane fusion protein”, Journal of virology, vol. 85, no. 13, pp. 6334–6342 doi:10.1128/JVI.00596-11 [doi].

